# Phasic arousal suppresses biases in mice and humans across domains of decision-making

**DOI:** 10.1101/447656

**Authors:** J. W. de Gee, K. Tsetsos, L. Schwabe, A.E. Urai, D. A. McCormick, M. J. McGinley, T. H. Donner

## Abstract

Decisions are often made by accumulating ambiguous evidence over time. The brain’s arousal systems are activated during such decisions. In previous work in humans, we showed that evoked responses of arousal centers during decisions are reported by rapid dilations of the pupil, and predict a suppression of biases in the accumulation of decision-relevant evidence (de Gee *et al.* 2017). Here, we show that this arousal-related suppression in decision bias acts on both conservative and liberal biases, and generalizes across species (humans / mice), sensory systems (visual / auditory), and domains of decision-making (perceptual / memory-based). In challenging sound-detection tasks, the impact of spontaneous or experimentally induced choice biases was reduced under high arousal. Similar bias suppression occurred when evidence was drawn from memory. All these behavioral effects were explained by reduced evidence accumulation biases. Our results pinpoint a general principle of the interplay between phasic arousal and decision-making.

## INTRODUCTION

The global arousal state of the brain changes from moment to moment (Aston-Jones & Cohen, 2005; McGinley, Vinck, et al., 2015). These global state changes are controlled in large part by modulatory neurotransmitters released from subcortical nuclei such as the noradrenergic locus coeruleus and the cholinergic basal forebrain. Release of these neuromodulators can profoundly change the operating mode of target cortical circuits (Aston-Jones & Cohen, 2005; Froemke, 2015; Harris & Thiele, 2011; Lee & Dan, 2012; Pfeffer et al., 2018). These same arousal systems are phasically recruited during elementary decisions, in relation to key computational variables such as uncertainty about making the correct choice and surprise about decision outcome (Aston-Jones & Cohen, 2005; Bouret & Sara, 2005; Colizoli, de Gee, Urai, & Donner, 2018; Dayan & Yu, 2006; Krishnamurthy, Nassar, Sarode, & Gold, 2017; Lak, Nomoto, Keramati, Sakagami, & Kepecs, 2017; Nassar et al., 2012; Parikh, Kozak, Martinez, & Sarter, 2007; Urai, Braun, & Donner, 2017).

Most decisions – including judgments about weak sensory signals in noise – are based on a protracted process of evidence accumulation (Shadlen & Kiani, 2013). This evidence accumulation process seems to be implemented in a distributed network of brain regions. In perceptual decisions, noise-corrupted decision evidence is encoded in sensory cortex, and downstream regions of association and motor cortices accumulate the fluctuating sensory response over time into a decision variable that forms the basis of behavioral choice (Bogacz, Brown, Moehlis, Holmes, & Cohen, 2006; Shadlen & Kiani, 2013; Siegel, Engel, & Donner, 2011; Wang, 2008). All these brain regions are impacted by the brain’s arousal system. Thus, arousal might alter the encoding of the momentary evidence, the accumulation thereof into a decision variable, and/or the implementation of the motor act.

We previously combined fMRI, pupillometry and behavioral modeling in humans to illuminate the interaction between phasic arousal and perceptual evidence accumulation (de Gee et al., 2017). We found that rapid pupil dilations during perceptual decisions report evoked responses in specific neuromodulatory (brainstem) nuclei controlling arousal, including the noradrenergic locus coeruleus (LC). We also showed that those same pupil responses predict a suppression of pre-existing biases in the accumulation of perceptual evidence. Specifically, in perceptual detection tasks, spontaneously emerging “conservative” biases (towards reporting the absence of a target signal) were reduced under large phasic arousal. Thus, it remains an open question whether phasic arousal stereotypically promotes liberal decision-making, or generally suppresses biases of any direction (conservative and liberal).

It is also unclear if the impact of arousal generalizes to higher-level, memory-based decisions. While elementary perceptual decisions are an established laboratory model of studying decision-making mechanisms, many important real-life decisions (e.g. which stock to buy) are also based on information gathered from memory. Recent advances indicate that memory-based decisions are also based on the accumulation of decision-relevant evidence; in this case, evidence drawn from memory (Shadlen & Shohamy, 2016).

Finally, it is also unknown if the impact of arousal on decision-making generalizes from humans to rodents. This is important because rodents are increasingly utilized as experimental models for decision-making mechanisms (Carandini & Churchland, 2013; Najafi & Churchland, 2018). Indeed, rodents (rats) can accumulate perceptual evidence in a similar fashion as humans (Brunton, Botvinick, & Brody, 2013), their arousal systems are homologously organized to those of humans (Amaral & Sinnamon, 1977; Berridge & Waterhouse, 2003), and pupil dilation reports arousal also in rodents (McGinley, David, & McCormick, 2015; Reimer et al., 2014; Vinck, Batista-Brito, Knoblich, & Cardin, 2015). But the interplay between phasic arousal and decision-making in rodents remains unknown.

Here, we address three key open issues pertaining to the interplay between arousal systems and decision-making: Does the dependence of decision biases on phasic arousal generalize (i) from humans to rodents, and (ii) from perceptual to memory-based decision-making? And (iii) does phasic arousal stereotypically promotes liberal decision-making, or reduce biases of any direction? We addressed these questions through a cross-species computational approach (Badre, Frank, & Moore, 2015), combining pupillometry and computational model-based analyses of behavior, in both humans and mice, and studying human decision-making in a variety of contexts.

## RESULTS

To address the role of phasic arousal in decision computations across species, we had humans and mice perform the same auditory go/no-go detection while measuring pupil-indexed phasic arousal. To test for additional generality, humans performed a forced choice decision task based on the same auditory evidence under systematic manipulations of target probabilities, and a memory-based decision task.

### In humans and mice, phasic arousal tracks a reduction of choice bias in an auditory detection task

We first trained mice (N = 5) and humans (N = 20) to report detection of a near-threshold auditory signal (Fig. 1A; Materials and Methods). Subjects searched for a signal (pure tone with a signal loudness that varied from trial to trial), which was embedded in a sequence of discrete, but dynamic, noise tokens (constant loudness). Because stable signals were embedded in fluctuating noise, detection performance could be maximized by accumulating the sensory evidence over time. To indicate a yes choice, mice licked for sugar water reward and human subjects pressed a button. Reaction times (RTs) decreased (Fig. 1B) and signal sensitivity (d’, from signal detection theory; Materials and Methods) increased (Fig. 1C) with signal strength (tone loudness; constant noise level). We quantified phasic arousal as the rising slope of the pupil, immediately after the onset of each sound token (hereafter called a “trial”). We chose this measure (i) for its temporal precision in tracking arousal during fast-paced tasks (Fig. 1D), (ii) to eliminate contamination by movements (see below, section *Suppression of bias by phasic arousal does not reflect motor preparation and is distinct from ongoing slow state fluctuations*), and (iii) to most specifically track noradrenergic activity (Reimer et al., 2016), which may play a specific role in decision-making (Aston-Jones & Cohen, 2005; Dayan & Yu, 2006). Because trial timing was predictable, subjects could anticipate the sound starts, and align their arousal response to trial onset. As such, pupil responses occurred as early as 40 ms after trial onset in mice (Fig. 1D), and from 240 ms after trial onset in humans (Fig. 1D). The shorter pupil response latencies in mice compared to humans might be due to their smaller eye and brain size.

**Figure 1.**
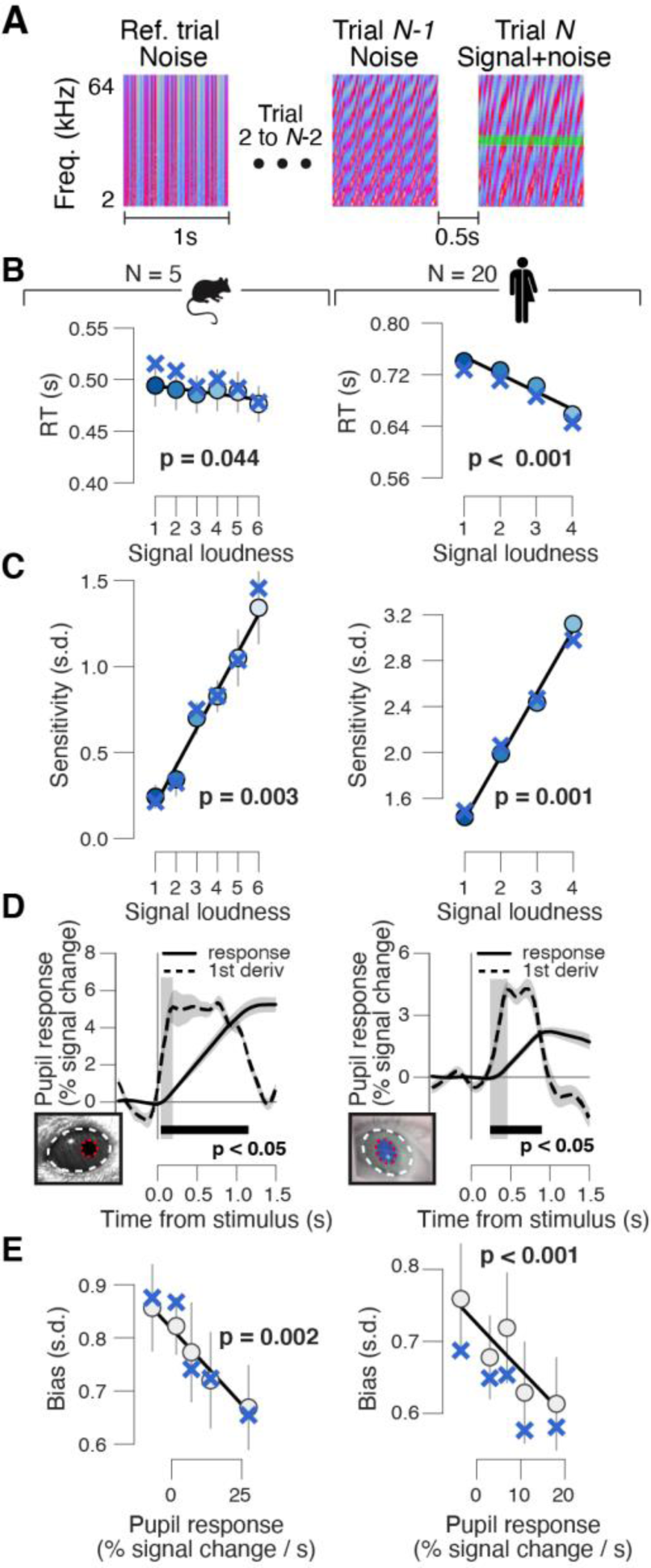
High phasic arousal is associated with reduced perceptual choice bias. **(A)** Auditory go/no-go tone-in-noise detection task. Schematic sequence of discrete sound tokens (trials) during a mini-block. Subjects were trained to respond to a weak signal (stable pure tone) in fluctuating noise and withheld a response for noise-only trials. Each sound token was treated as a separate decision (Materials and Methods). **(B)** Relationship between reaction time and signal loudness in mice (left) and humans (right). ‘X’ symbols are predictions from best fitting variant of drift diffusion model (Materials and Methods); stats are from mixed linear modeling (Materials and Methods). **(C)** As panel B, but for sensitivity (quantified by d’ from signal detection theory; Materials and Methods**). (D)** Task-evoked pupil response (solid line) and response derivative (dashed line) in mice (left) and humans (right). Grey window, interval for task-evoked pupil response measures (Materials and Methods); black bar, significant pupil derivative; stats, cluster-corrected one-sample t-test. **(E)** Relationship between overall perceptual choice bias (Materials and Methods) and task-evoked pupil response in mice (left) and humans (right). Subjects likely set only a single decision criterion because signal strength was drawn pseudo-randomly on each trial (see also Fig. S1H,M). Linear fits were plotted if first-order fit was superior to constant fit; quadratic fits were not superior to first-order fits (Materials and Methods). ‘X’ symbols are predictions from best fitting variant of drift diffusion model (Materials and Methods); stats, mixed linear modeling. Panels B-E: group average (N = 5; N = 20); shading or error bars, s.e.m.

Pupil responses occurred on all trials, whether or not there was a behavioral response (Fig. S1E,J). However, as in our earlier work (de Gee et al., 2017), we found a consistent relationship between the early, task-evoked pupil response and choice bias, in mice and humans (Fig. 1E). Both species had an overall conservative choice bias, often failing to report the signal tones (Fig. 1E). This bias was partially suppressed on trials with large pupil responses (Fig. 1E, computed after collapsing across signal strengths; see also Fig. S1C and Materials and Methods). Unlike the consistent effect on bias, phasic pupil responses exhibited a less consistent relationship to d’ and RT (Fig. S1G,L).

Previous work has associated baseline, pre-stimulus arousal state with non-monotonic (inverted U-shape) effects on decision performance (Aston-Jones & Cohen, 2005; Yerkes & Dodson, 1908), including in the same mouse dataset analyzed here (McGinley, David, et al., 2015). Unlike baseline arousal, and in line with earlier observations (de Gee et al., 2017), we here found that pupil-linked phasic (task-evoked) arousal had a monotonic (linear) effect on bias (Fig. 1E; Materials and Methods), pointing to distinct functional roles of tonic and phasic arousal, perhaps resulting from the distinct neuromodulators or modulatory receptors (Reimer et al., 2016).

### In humans and mice, pupil-linked bias reduction is in the sensory evidence accumulation process

Fitting decision-making models to choices and reaction times enabled us to gain deeper insight into how the decision process was affected by phasic pupil-linked arousal. We fitted the drift diffusion model (Fig. S2A), which is a widely used bounded accumulation model of decision-making (Bogacz et al., 2006; Brody & Hanks, 2016; Gold & Shadlen, 2007; Ratcliff & McKoon, 2008), a class of models that describe the accumulation of noisy sensory evidence in a decision variable that drifts to one of two bounds. We used the diffusion model to quantify the effects of pupil-linked arousal on the following components of the decision process: the starting point of evidence accumulation, the evidence accumulation itself (the mean drift rate and an evidence-dependent bias in the drift, henceforth called “drift bias”), boundary separation (implementing speed-accuracy tradeoff) and the so-called non-decision time (the speed of pre-decisional evidence encoding and post-decisional translation of choice into motor response). We allowed all parameters except for starting point to vary with pupil response amplitude (see Materials and Methods for justification). The model accounted well for the overall behavior, revealing the expected increase of drift rate with signal strength (Fig. S2D,G) and accurate prediction of RTs and sensitivity (blue ‘X’ markers in Fig. 1B,C).

In both species, the starting point was biased towards no-go (Fig. S2C,F). Overcoming this conservative bias set by starting point required shifting the drift bias towards the yes bound, which occurred on trials with large pupil responses (Fig. 2C). The relationship between pupil responses and drift bias was linear (Fig. 2C), and accurately predicted the pupil-dependent reduction in overt conservative choice bias (blue ‘X’ markers in Fig. 1E).

**Figure 2.**
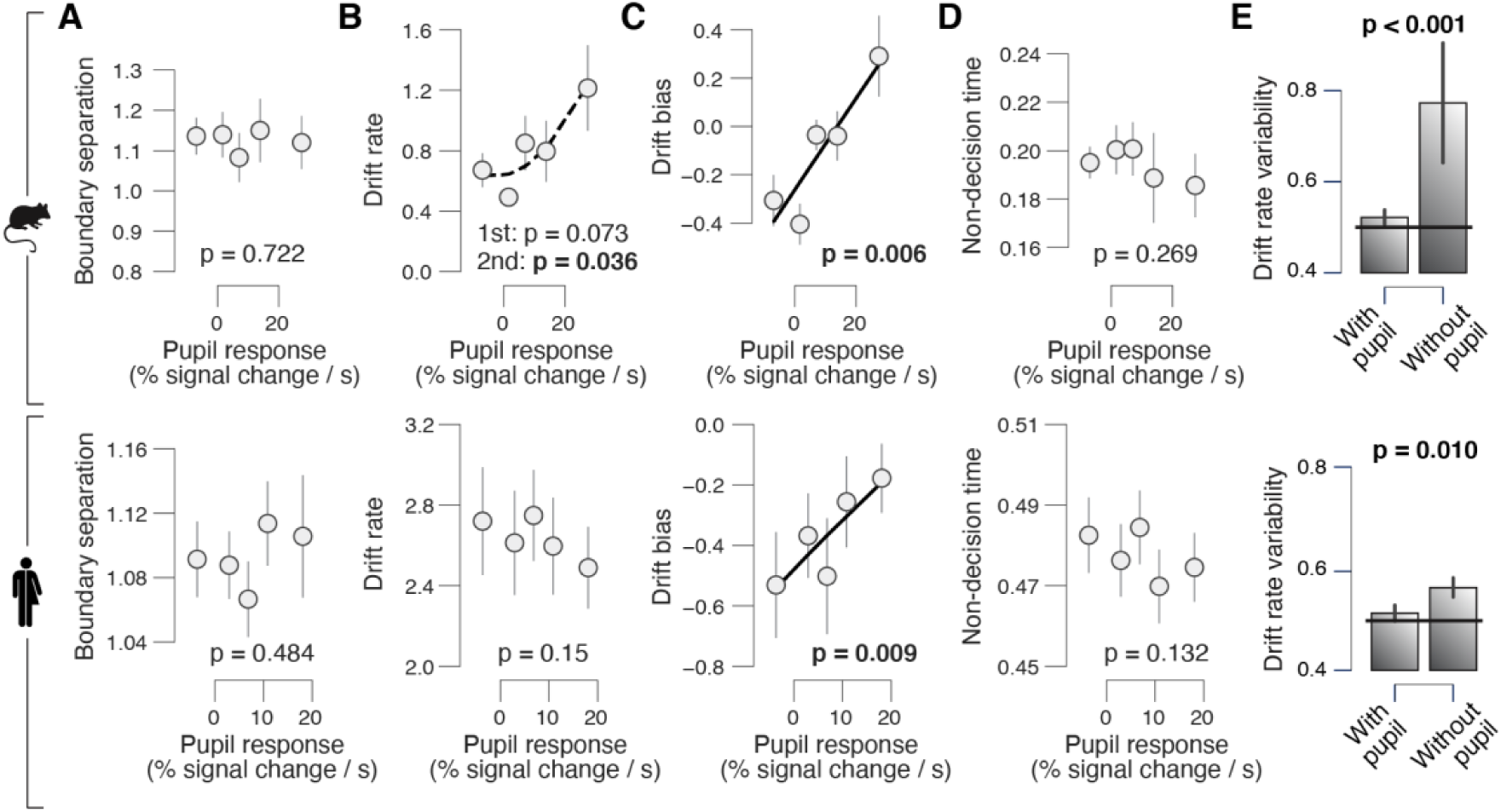
Phasic arousal reduces overt choice bias by reducing a bias in evidence accumulation. **(A)** Relationship between boundary separation estimates and task-evoked pupil response in mice (top) and humans (bottom), collapsed across signal loudness. See Fig. S2 for parameter estimates separately per signal loudness. Linear fits are plotted wherever the first-order fit was superior to the constant fit; quadratic fits were plotted (dashed lines) wherever the second-order fit was superior to first-order fit. Stats, mixed linear modeling. **(B-D)** As A, but for drift rate, drift bias and non-decision time estimates, respectively. **(E)** Recovered drift rate variability for models with and without pupil predicted shift in drift bias. The model was fit to simulated RT distributions from two conditions that differed according to the fitted drift bias estimates in the lowest and highest pupil-defined bin of each individual (Materials and Methods). Black line, true (simulated) drift rate variability. Stats, paired-samples t-test. All panels: group average (N = 5; N = 20); error bars, s.e.m.

Phasic arousal had no, or less consistent, effects on the other model parameters. There was no consistent monotonic effect of pupil response on boundary separation, drift rate or non-decision time in either species (mice: p = 0.722, p = 0.073 and p = 0.269, respectively; humans: p = 0.484, p = 0.15 and p = 0.132, respectively; Fig. 2). Without collapsing across signal loudness we observed a positive (negative) relationship between drift rate and pupil response in mice (humans) (Fig. S2D,G), and a negative relationship between pupil response and non-decision time in both mice and humans (Fig. S2D,G). Thus, the major impact of phasic arousal in this task was to suppress a bias in evidence accumulation.

Perceptual choice variability has been attributed to evidence accumulation noise, rather than to systematic accumulation biases, under the assumption that biases will remain constant across trials (Drugowitsch, Wyart, Devauchelle, & Koechlin, 2016). Instead, our results show that accumulation biases vary dynamically across trials as a function of phasic arousal. This indicates that the resulting choice variations should appear as random trial-by-trial variability in evidence accumulation when ignoring phasic arousal. We found that this was the case in our data (Fig. 2E). We simulated RT distributions from two conditions that differed according to the fitted drift bias estimates in the lowest and highest pupil-defined bin of each individual (Materials and Methods). The diffusion model accounts for trial-to-trial accumulation “noise” with the drift rate variability parameter (Bogacz et al., 2006; Ratcliff & McKoon, 2008). When fitting the model to these simulated RT distributions, drift rate variability was accurately recovered when drift bias could vary with condition but was overestimated when drift bias was fixed (Fig. 2E). Note that this analysis is agnostic about the source of trial-by-trial variations in phasic arousal, which was not under experimental control in the present study (but see (Colizoli et al., 2018; Nassar et al., 2012; Urai et al., 2017). But the results clearly show that a significant fraction of choice variability does not originate from noise within the evidence accumulation machinery itself.

### Phasic arousal predicts a reduction of conservative and liberal biases in perceptual evidence accumulation

The majority of mice and humans in the above go/no-go task exhibited a conservative bias, or tendency towards choosing “no”, which was reduced on trials with large task-evoked pupil responses. We thus wondered whether phasic arousal promotes liberal decision-making, irrespective of overall bias, or rather whether it suppresses both liberal and conservative biases.

To arbitrate between these scenarios, we asked a new group of humans subjects (N=15) to perform a forced choice (yes/no) version of the task, using the same type of auditory evidence (Fig. 3A), but with different probability of signal occurrence in blocks. In “rare” blocks, the signal occurred in 30% of trials, and in “frequent” blocks, the signal occurred in 70% of trials (Fig. 3B; Material and Methods). As expected (Green & Swets, 1966), subjects developed a conservative bias in the rare signal condition and a liberal bias in the frequent signal condition (Fig. 3D,G). Phasic-pupil responses predicted a change in choice biases towards neutrality for both block types (which occurred within the same experimental session) (Fig. 3E,H). As observed in the go/no-go task (Fig. 2), the effect of pupil-linked arousal on perceptual choice biases was mediated by shifts in accumulation biases (Fig. 3F,I). There was an effect of pupil-linked arousal on starting point too, but in the opposite direction as the perceptual choice bias shift (Fig. S3D,G). The pupil-linked changes in drift bias, but less so the changes in starting point, correlated with the individual reductions in perceptual choice bias as measured by SDT in the rare condition (squared multiple correlation R2 = 0.959; drift bias: beta = -0.97, p < 0.001; starting point: beta = -0.07, p = 0.039) as well as in the frequent condition (squared multiple correlation R2 = 0.997; drift bias: beta = -1.08, p < 0.001; starting point: beta = 0.29, p = 0.024). Thus, only the changes of drift bias explained the reductions in perceptual choice bias. Pupil responses were not associated with RT or sensitivity, nor with the drift diffusion model parameters boundary separation, drift rate or non-decision time (Fig. S3B–G). In sum, phasic arousal reduces choice biases irrespective of direction (conservative or liberal).

**Figure 3.**
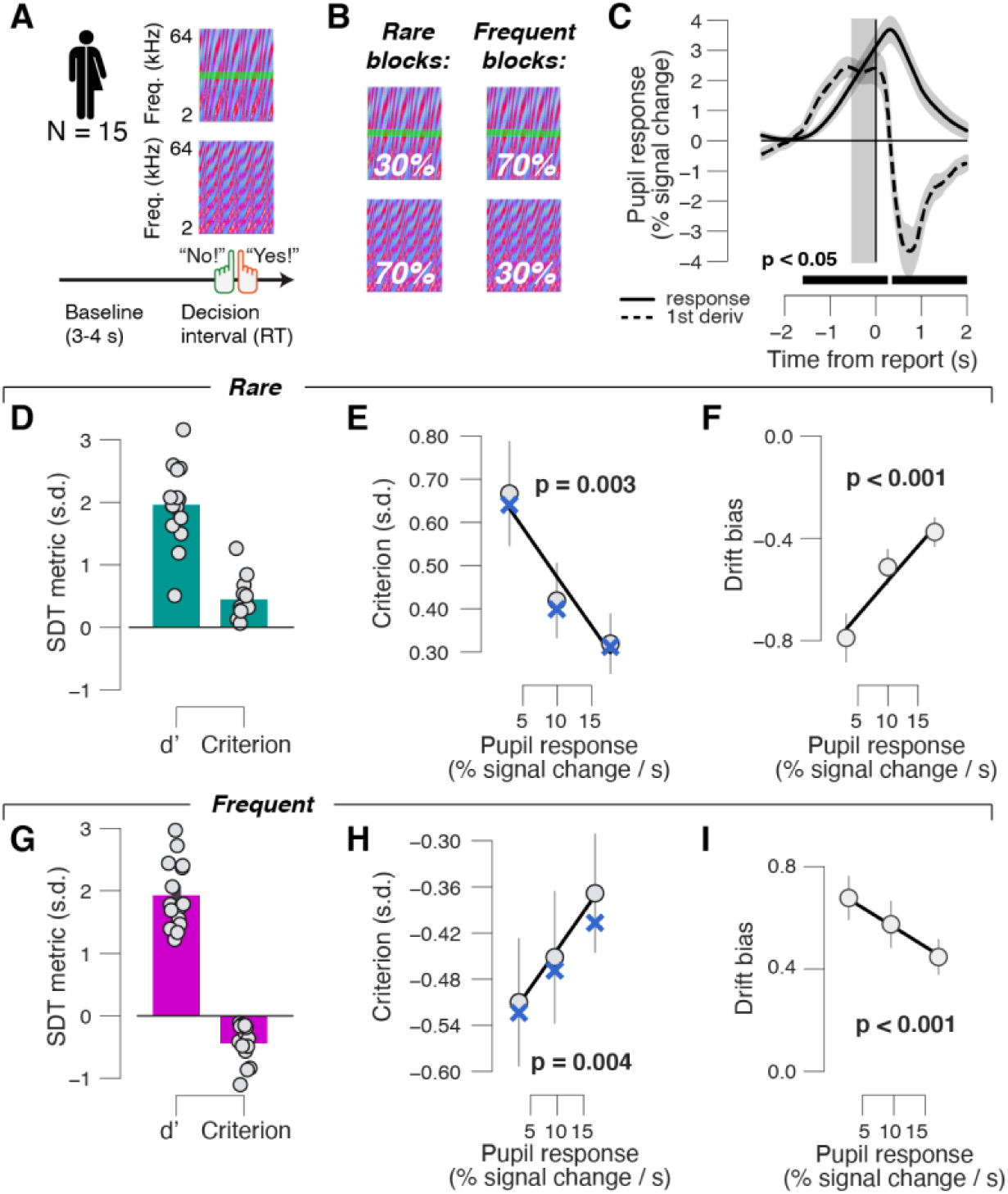
Phasic arousal reduces both conservative and liberal accumulation biases. **(A)** Auditory yes/no (forced choice) tone-in-noise detection task. Schematic sequence of events during a trial. Subjects reported the presence or absence of a faint signal (pure sine wave) embedded in noise (Materials and Methods). **(B)** Signal occurrence was varied across blocks of trials. **(C)** Task-evoked pupil response (solid line) and response derivative (dashed line). Grey, interval for task-evoked pupil response measures (Materials and Methods); black bar, significant pupil derivative. Stats, paired-samples t-test. **(D)** Overall sensitivity and choice bias (quantified by signal detection d’ and criterion, respectively) in the rare signal condition (P(Signal) = 0.3). **(E)** Relationship between choice bias and task-evoked pupil response in the rare condition. Linear fits were plotted if first-order fit was superior to constant fit; quadratic fits were not superior to first-order fits. ‘X’ symbols are predictions from the drift diffusion model; stats, mixed linear modeling. We used three pupil-defined bins because there were fewer critical trials per individual (less than 500) than in the previous data sets (more than 500; Materials and Methods). **(F)** As E, but for relationship between drift diffusion model parameter drift bias and task-evoked pupil response. **(G-I)** As D-F, but for the frequent signal condition (P(Signal) = 0.7). All panels: group average (N = 15); error bars, s.e.m.

### Phasic arousal predicts reduction of accumulation biases in memory-based decisions

Many decisions are not based on current sensory signals, but on information drawn from memory. It has been proposed that memory-based decisions follow the same sequential sample principle established for perceptual decisions, whereby the “samples” accumulated into the decision variable are drawn from memory (Bowen, Spaniol, Patel, & Voss, 2016; Ratcliff, 1978; Shadlen & Shohamy, 2016). We thus assessed whether the arousal-dependent bias suppression identified for perceptual decisions, above, generalized to memory-based decisions.

We modeled the impact of pupil-linked phasic arousal on choice behavior in a yes/no recognition memory task (Fig. 4A; Materials and Methods). Subjects (N=54) were instructed to memorize 150 pictures (intentional encoding) and to evaluate how emotional each picture was on a 4-point scale from 0 (“neutral”) to 3 (“very negative”). Twenty-four hours post encoding, subjects saw all pictures that were presented on the first day and an equal number of novel pictures in randomized order, and indicated for each item whether it had been presented before (“yes – old”) or not (“no – new”). We observed that subject-wise overall biases (irrespective of pupil responses) varied substantially across the 54 individuals, from strongly liberal to strongly conservative (Fig. 4B, x-axis and colors). Therefore, in addition to probing for memory-based decisions, this data set afforded a subject-wise test of the direction-dependence (conservative vs. liberal) of the pupil-linked arousal effect.

**Figure 4.**
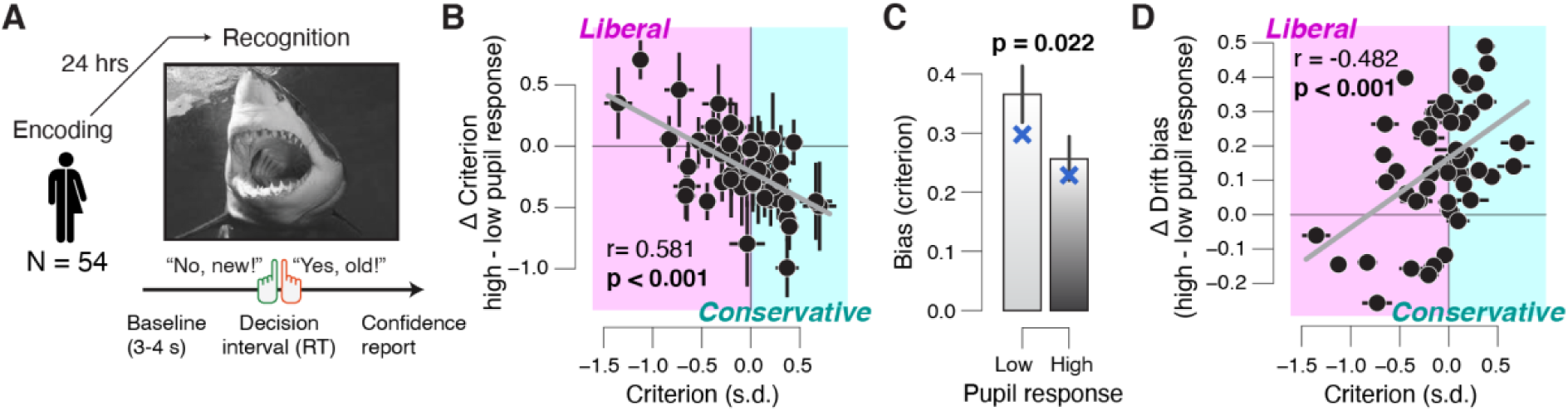
Phasic arousal predicts a reduction of memory accumulation biases. **(A)** A Picture yes/no (forced choice) recognition task. Schematic sequence of events during a trial. Subjects judged whether they had seen pictures twenty-four hours previously during an encoding task (Materials and Methods). See Fig. S5A,B for task-evoked pupil response time course. **(B)** Individual pupil predicted shift in choice bias (quantified by signal detection criterion), plotted against individual’s overall choice bias. Data points, individual subjects. Stats, Pearson’s correlation. Error bars, 60% confidence intervals (bootstrap). **(C)** Choice bias (sign-flipped for overall liberal subjects) for low and high pupil response bins. ‘X’ symbols are predictions from the drift diffusion model; stats, paired-samples t-test; group average (N = 54); error bars, s.e.m. **(D)** As B, but pupil predicted shift in drift bias, plotted against individual’s overall choice bias.

We observed a robust relationship between subjects’ overall choice bias, and the pupil predicted shift in that bias. Namely, those subjects with the strongest biases, whether liberal or conservative, exhibited the strongest pupil-predicted shift towards a neutral bias (Fig. 4B). The group-average choice bias (signal detection theoretic criterion; sign-flipped for overall-liberal subjects) was significantly reduced towards 0 on large pupil response trials (Fig. 4C). Again, this behavioral effect was explained by pupil-linked changes in drift bias (Fig. 4D, Fig. S4E), not by changes in starting point bias (Fig. S4E,F) (difference in correlation: Δr = -0.599, p < 0.001). Note that lower criterion values indicate more liberal behavior, and lower drift bias values more conservative behavior, which explains the correlations of opposite sign in Fig. 4 panels B and D. The pupil response further predicted an increase in drift rate (p = 0.012), a lengthening of non-decision time (p=0.017), and no changes in starting point (p = 0.161) or boundary separation (p = 0.089) (Fig. S4E).

We conclude that, similar to perceptual choices, phasic arousal reduces both liberal and conservative memory-based choice biases.

### Suppression of bias by phasic arousal does not reflect motor preparation and is distinct from ongoing slow state fluctuations

One concern in the go/no-go task is that the bias suppression associated with large pupil responses was produced by the motor response for go choices. Specifically, the central input to the pupil contains a sustained component during evidence accumulation, followed by a transient at the motor response (de Gee et al., 2017; de Gee, Knapen, & Donner, 2014; Hupé, Lamirel, & Lorenceau, 2009; Murphy, Boonstra, & Nieuwenhuis, 2016). The sustained component might entail motor preparatory activity (Donner et al, 2009). Thus, these components (motor preparatory activity and transient activity at lick / button-press) could have contributed to the pupil response amplitudes on go trials, but not (or less so) on no-go trials. This motor component could theoretically produce the observed relationship between pupil responses and choice bias.

We reduced contamination by the transient motor-related component by focusing on the early component of pupil dilation (Materials and Methods). However, it is possible that this did not fully correct for the asymmetry between go and no-go trials in motor preparatory activity. To further address this concern, we re-analyzed results from the forced choice (yes/no) version of the task, which were published previously (de Gee et al., 2017). In this task, motor responses, and thus likely associated preparatory activity, were balanced across yes and no choices (Fig. 3A; Materials and Methods). Importantly, pupil response amplitudes in the go/no-go and yes/no tasks were correlated across eighteen human subjects who participated in both experiments (Fig. S5C). This was true for both yes choices and no choices. As ‘no’ choices were not matched in terms of motor responses (no-go vs. button press), this result indicates that pupil response amplitudes primarily reflected decision processing rather than motor preparation. Furthermore, consistent with our go/no-go results in mice and humans, we observed an arousal-linked suppression of perceptual choice bias (Fig. S5D) and suppression of evidence accumulation bias (Fig. S5F). Taken together, we conclude that the suppression of choice bias in our results does not reflect motor preparation.

A second concern is that the bias suppression effects under large pupil responses reported here might be “inherited” through a previously observed negative correlation between phasic pupil responses and pre-trial baseline pupil diameter (de Gee et al., 2014; Gilzenrat, Nieuwenhuis, Jepma, & Cohen, 2010). Variation in pre-trial pupil size causes floor and ceiling effects on phasic dilations, shaped by light conditions. A dependence on baseline pupil could not account for the results reported here, for four reasons. First, in the go/no-go data sets, pupil responses were quantified as the rising slopes (see above), and those exhibited a negligible correlation to the preceding baseline diameter (mice: r = 0.014 ±0.028 s.e.m.; humans: r = -0.037 ±0.017 s.e.m.). Second, there was a non-monotonic association between baseline pupil diameter and decision bias in mice (McGinley, David, et al., 2015), in contrast to the monotonic pattern we observed here for phasic arousal in the same dataset (Fig. 1E). Third, although the pupil responses were negatively related to baseline pupil diameter in the yes/no rare condition (r = - 0.163, ±0.041 s.e.m.) and the yes/no frequent condition (r = -0.109, ±0.047 s.e.m.), there was either no, or a weak, systematic association between baseline pupil diameter and decision bias (yes/no rare: p = 0.043; yes/no frequent: p = 0.556). Fourth, in the yes/no recognition task, there was again a negligible correlation between pupil response and preceding baseline diameter (r = -0.010 ±0.010 s.e.m.). Thus, the behavioral correlates of pupil responses reported in this paper reflect genuine effects of phasic arousal, which are largely uncontaminated by the baseline, slowly varying, arousal state.

## DISCUSSION

Arousal is traditionally thought to globally upregulate the efficiency of information processing (e.g., the quality of evidence encoding or the efficiency of accumulation (Aston-Jones & Cohen, 2005; McGinley, Vinck, et al., 2015)). However, recent work indicates that phasic arousal signals might have distinct effects, such as reducing the impact of prior expectations and biases on decision formation (de Gee et al., 2017, 2014; Krishnamurthy et al., 2017; Nassar et al., 2012; Urai et al., 2017). We here established a principle of the function of phasic arousal in decision-making, which generalizes across species (humans and mice) and domains of decision-making (from perceptual to memory-based): phasic arousal suppresses biases in the accumulation of evidence leading up to a choice.

We observed the bias-suppression principle in human and mouse choice behavior during analogous auditory tone-detection tasks. Task-evoked pupil responses occurred early during decision formation, even on trials without a motor response, and predicted a suppression of conservative choice bias. Behavioral modeling revealed that the bias reduction was due to a selective interaction with the accumulation of information from the noisy sensory evidence. We further showed that phasic arousal reduces accumulation biases, whether conservative or liberal, as seen in the conditions with different stimulus presentation statistics. Finally, we established that pupil-linked suppression of evidence accumulation bias also occurs for memory-based decisions. We conclude that the ongoing deliberation, culminating in a choice (Shadlen & Kiani, 2013), is shaped in a canonical way by transient boosts in the global arousal state of the brain: suppression of evidence accumulation bias.

We here used pupil responses as a peripheral readout of changes in cortical arousal state (Larsen & Waters, 2018; McGinley, Vinck, et al., 2015). Indeed, recent work has shown that pupil diameter closely tracks several established measures of cortical arousal state (Larsen & Waters, 2018; McGinley, David, et al., 2015; McGinley, Vinck, et al., 2015; Reimer et al., 2014; Vinck et al., 2015). Changes in pupil diameter have been associated with locus coeruleus (LC) activity in humans (de Gee et al., 2017; Murphy, O’Connell, O’Sullivan, Robertson, & Balsters, 2014), monkeys (Joshi, Li, Kalwani, & Gold, 2016; Varazzani, San-Galli, Gilardeau, & Bouret, 2015), and mice (Breton-Provencher & Sur, 2019; Liu, Rodenkirch, Moskowitz, Schriver, & Wang, 2017; Reimer et al., 2016). However, some of these studies also found unique contributions to pupil size in other subcortical regions, such as the cholinergic basal forebrain and dopaminergic midbrain, and the superior and inferior colliculi (de Gee et al., 2017; Joshi et al., 2016; Reimer et al., 2016). Thus, the exact neuroanatomical and neurochemical source(s), of our observed effects of phasic arousal on decision-making, remain to be determined.

There is mounting evidence for an arousal-linked reduction of biases (or priors) in humans, and our current findings generalize this emerging principle to rodents (mice) as well as to a higher-level form of evidence accumulation: memory-based. Previous work has shown a similar suppression of evidence accumulation bias during challenging visual perceptual choice tasks (contrast detection and random dot motion discrimination) (de Gee et al., 2017). Similarly, during sound localization in a dynamic environment, phasic arousal predicts a reduced influence of prior expectations on perception (Krishnamurthy et al., 2017). Furthermore, suppressive effects of phasic arousal also apply to choice history biases that evolve across trials. In this case, phasic arousal reflects perceptual decision uncertainty on the current trial and a reduction of choice repetition bias on the next trial (Urai et al., 2017). One study reported that phasic arousal predicted an overall reduction of reaction time during random dot motion discrimination (van Kempen et al., 2019). However, in this study, the signal strength was so high that the task could be solved without the temporal accumulation of evidence: reaction times were overall short (median < 600 ms) and performance was close to ceiling. Thus, the arousal-dependent bias suppression seems to be a principle that generalizes across species, directions of bias, and domains of decision-making.

We chose the drift diffusion model to capture our behavioral data because the model: (i) is sufficiently low-dimensional so that its parameter estimates are well constrained by the choices and shape of RT distributions, (ii) has been shown to successfully account for behavioral data from a wide range of decision-making tasks, including go/no-go (Ratcliff, Huang-Pollock, & McKoon, 2016; Ratcliff & McKoon, 2008), and (iii) is, under certain parameter regimens, equivalent to a parameter reduction of biophysically detailed neural circuit models of decision-making (Bogacz et al., 2006; Wong & Wang, 2006). The drift diffusion model relies on three main assumptions. First, in the go/no-go task, we assume that participants accumulated the auditory evidence *within* each trial (discrete noise token) during a mini-block and reset this accumulation process before the next discrete sound. Second, in both the go/no-go and yes/no tasks, we assume that subjects actively accumulated evidence towards both yes and no choices, which is supported by neurophysiological data from yes/no tasks (Deco, Pérez-Sanagustín, de Lafuente, & Romo, 2007; Donner, Siegel, Fries, & Engel, 2009). Third, in the go/no-go task, we assume that subjects set an implicit boundary for no choices (Ratcliff et al., 2016). The quality of our model fits suggest that the model successfully accounted for the measured behavior, lending support to the validity of these assumptions.

The monotonic effects of “phasic” arousal on decision biases that we report here contrast with observed effects of “tonic” (pre-stimulus) arousal, which has a non-monotonic (inverted U) effect on behavior (perceptual sensitivity and bias) and neural activity (the signal-to-noise ratio of thalamic and cortical sensory responses) (Gelbard-Sagiv, Magidov, Sharon, Hendler, & Nir, 2018; McGinley, David, et al., 2015). Our study allows a direct comparison of tonic and phasic arousal effects within the same data set in mice. A previous report on that data set showed that the mice’s behavioral performance was most rapid, accurate, and the least biased at intermediate levels of ‘tonic’ arousal (medium baseline pupil size) (McGinley, David, et al., 2015). In contrast, we here show that their behavioral performance was linearly related to phasic arousal, with the most rapid, accurate and least biased choices for the largest phasic arousal responses. It is tempting to speculate that these differences result from different neuromodulatory systems governing tonic and phasic arousal. Indeed, rapid dilations of the pupil (phasic arousal) are more tightly associated with phasic activity in noradrenergic axons, whereas slow changes in pupil (tonic arousal) are accompanied by sustained activity in cholinergic axons (Reimer et al., 2016). However, more work is needed to directly relate phasic and tonic pupil dilation during decision-making to neuromodulatory systems in the brain.

Recent findings indicate that intrinsic behavioral variability is increased during sustained (“tonic”) elevation of NA levels, in line with the “adaptive gain theory” (Aston-Jones & Cohen, 2005). First, optical stimulation of LC inputs to anterior cingulate cortex caused rats to abandon strategic counter prediction in favor of stochastic choice in a competitive game (Tervo et al., 2014). Second, chemogenetic stimulation of the LC in rats performing a patch leaving task increased decision noise and subsequent exploration (Kane et al., 2017). Third, pharmacologically reducing central noradrenaline levels in monkeys performing an operant effort exertion task parametrically increased choice consistency (Jahn et al., 2018). Finally, pharmacologically increasing central tonic noradrenaline levels in human subjects boosted the rate of alternations in a bistable visual input and long-range correlations in brain activity (Pfeffer et al., 2018). Here, we tested for the effect of phasic arousal on a range of behavioral parameters, including decision noise. In the drift diffusion model, increased decision noise would manifest as a decrease of the mean drift rate, which scales inversely with (within-trial) decision noise. We found no such effect that was consistent across data sets. This is another indication, together with the baseline pupil effects reported by (McGinley, David, et al., 2015), that the effects of phasic and tonic neuromodulation are distinct.

One influential account holds that phasic LC responses during decision-making are triggered by the threshold crossing in some circuit accumulating evidence, and that the resulting noradrenaline release then facilitates the translation of the choice into a motor act (Aston-Jones & Cohen, 2005). Within the drift diffusion model, this predicts a reduction in non-decision time and no effect on evidence accumulation. In contrast to this prediction, we found that in all our datasets that phasic arousal affected evidence accumulation (suppressing biased therein), but not non-decision time. Our approach does not enable us to rule out an effect of phasic arousal on movement execution (i.e., kinematics). Yet, our results clearly establish an important role of phasic arousal in evidence accumulation, ruling out any *purely* post-decisional account. This implies that phasic LC responses driving pupil dilation are already recruited during evidence accumulation, or that the effect of pupil-linked arousal on evidence accumulation are governed by systems other than the LC. Future experiments characterizing phasic activity in the LC or other brainstem nuclei involved in arousal during protracted evidence accumulation tasks could shed light on this issue.

Anatomical evidence supports the speculation that task-evoked neuromodulatory responses and cortical decision circuits interact in a recurrent fashion. One possibility is that neuromodulatory responses alter the balance between “bottom-up” and “top-down” signaling across the cortical hierarchy (Friston, 2010; Hasselmo, 2006; Hsieh, Cruikshank, & Metherate, 2000; Kimura, Fukuda, & Tsumoto, 1999; Kobayashi et al., 2000). Sensory cortical regions encode likelihood signals and sent these (bottom-up) to association cortex; participants’ prior beliefs (about for example target probability) are sent back (top-down) to the lower levels of the hierarchy (Beck, Ma, Pitkow, Latham, & Pouget, 2012; Pouget, Beck, Ma, & Latham, 2013). Neuromodulators might reduce the weight of this prior in the inference process (Friston, 2010; Moran et al., 2013), thereby reducing choice biases. Another possibility is neuromodulator release might scale with uncertainty about the incoming sensory data (Friston, 2010; Moran et al., 2013). Such a process could be implemented by top-down control of the cortical systems decision-making over neuromodulatory brainstem centers. This line of reasoning is consistent with anatomical connectivity (Aston-Jones & Cohen, 2005; Sara, 2009). Finally, a related conceptual model that has been proposed for phasic LC responses is that cortical regions driving the LC (e.g. ACC) continuously compute the ratio of the posterior probability of the state of the world, divided by its (estimated) prior probability (Dayan & Yu, 2006). LC is then activated when neural activity ramps towards the non-default choice (against ones’ bias). The resulting LC activity might reset its cortical target circuits (Bouret & Sara, 2005) and override the default state (Dayan & Yu, 2006), facilitating the transition of the cortical decision circuitry towards the non-default state. These scenarios are in line with recent insights that (LC-mediated) pupil-linked phasic arousal shapes brain-wide connectivity (Stitt, Zhou, Radtke-Schuller, & Fröhlich, 2018; Zerbi et al., 2019).

Our study showcases the value of comparative experiments in humans and non-human species. One would expect the basic functions of arousal systems (e.g. the LC-NA system) to be analogous in humans and rodents. Yet, it has been unclear whether these systems are recruited in the same way during decision-making. Computational variables like decision uncertainty or surprise are encoded in prefrontal cortical regions (e.g. anterior cingulate or orbitofrontal cortex (Kepecs, Uchida, Zariwala, & Mainen, 2008; Ma & Jazayeri, 2014; Pouget, Drugowitsch, & Kepecs, 2016) and conveyed to brainstem arousal systems via top-down projections (Aston-Jones & Cohen, 2005; Breton-Provencher & Sur, 2019). Both the cortical representations of computational variables and top-down projections to brainstem may differ between species. More importantly, it has been unknown whether key components of the decision formation process, in particular evidence accumulation, would be affected by arousal signals in the same way between species. Only recently has it been established that rodents (rats) and humans accumulate perceptual evidence in an analogous fashion (Brunton et al., 2013). Here, we established that the shaping of evidence accumulation by phasic arousal is also governed by a conserved principle.

## MATERIALS AND METHODS

### Subjects

All procedures concerning the animal experiments were carried out in accordance with Yale University Institutional Animal Care and Use Committee, and are described in detail elsewhere (McGinley, David, et al., 2015). Human subjects were recruited and participated in the experiment in accordance with the ethics committee of the Department of Psychology at the University of Amsterdam (go/no-go and yes/no task), the ethics committee of Baylor College of Medicine (yes/no task with biased signal probabilities) or the ethics committee of the University of Hamburg (recognition task). Human subjects gave written informed consent and received credit points (go/no-go and yes/no tasks) or a performance-dependent monetary remuneration (yes/no task with biased signal probabilities and recognition task) for their participation. We analyzed two previously unpublished human data sets, and re-analyzed a previously published mice data set (McGinley, David, et al., 2015) and two human data sets (Bergt, Urai, Donner, & Schwabe, 2018; de Gee et al., 2017). Bergt *et al.* (2018) have analyzed pupil responses only during the encoding phase of the recognition memory experiment; we here present the first analyses of pupil responses during the recognition phase. We selected the data set by Bergt *et al* (2018) for across-subject correlations of variables of interest (Fig. 4). This data set had a sufficient sample size for such an analysis, based on the effect size obtained in a previous study (de Gee et al., 2014). The sample sizes (and trial numbers per individual) for the newly collected human data sets were determined based on the effects observed in a previous study comparing diffusion model parameters between pupil conditions: (de Gee et al., 2017) with N=14.

Five mice (all males; age range, 2–4 months) and twenty human subjects (15 females; age range, 19– 28 y) performed the go/no-go task. Twenty-four human subjects (of which 18 had already participated in the go/no-go task; 20 females; age range, 19–28 y) performed an additional yes/no task. Fifteen human subjects (8 females; age range, 20–28 y) performed the yes/no task with biased signal probabilities. Fifty-four human subjects (27 females; age range, 18–35 y) performed a picture recognition task, of which two were excluded from the analyses due to eye-tracking failure.

For the go/no-go task, mice performed between five and seven sessions (described in (McGinley, David, et al., 2015)), yielding a total of 2469–3479 trials per subject. For the go/no-go task, human participants performed 11 blocks of 60 trials each (distributed over two measurement sessions), yielding a total of 660 trials per participant. For the yes/no task, human participants performed between 11 and 13 blocks of 120 trials each (distributed over two measurement sessions), yielding a total of 1320–1560 trials per participant. For the yes/no task with biased signal probabilities, human subjects performed 8 blocks of 120 trials each (distributed over two measurement sessions), yielding a total of 960 trials per participant. For the picture recognition task, human subjects performed 300 trials.

### Behavioral tasks

#### Perceptual (auditory tone-in-noise) go/no-go detection task

Each mini-block consisted of two to seven consecutive trials. Each trial was a distinct auditory noise stimulus of 1 s duration, and the inter-trial interval was 0.5 s. A weak signal tone was added to the last trial in the mini-block (Fig. 1A). The number of trials, and thus the signal position in the sequence, was randomly drawn beforehand. The probability of a target signal decreased linearly with trial number (Fig. S1A), so as to flatten the hazard rate of signal onset across the mini-block. Each mini-block was terminated by the subject’s go response (hit or false alarm) or after a no-go error (miss). Each trial consisted of an auditory noise stimulus, or a pure sine wave added to one of the noise stimuli (cosine-gated 2 kHz for humans; new tone frequency each session for mice to avoid cortical reorganization across the weeks of training (McGinley, David, et al., 2015)). Noise stimuli were temporally orthogonal ripple combinations, which have spectro-temporal content that is highly dynamic, thus requiring temporal integration of the acoustic information in order to detect the stable signal tones (McGinley, David, et al., 2015). In the mouse experiments, auditory stimuli were presented at an overall intensity of 55dB (root-mean-square [RMS] for each 1 second trial). In the human experiments, auditory stimuli were presented at an intensity of 65dB using an IMG Stageline MD-5000DR over-ear headphone.

Mice learned to respond during the signal-plus-noise trials and to withhold responses during noise trials across training sessions. Mice responded by licking for sugar water reward. Human participants were instructed to press a button with their right index finger. Correct yes choices (hits) were followed by positive feedback: 4 μL of sugar water in the mice experiment, and a green fixation dot in the human experiment. In both mice and humans, false alarms were followed by an 8 s timeout. Humans, but not mice, also received an 8 s timeout after misses. This design difference was introduced to compensate for differences in general response bias between species evident in pilot experiments: while mice tended to lick too frequently without a selective penalty for false alarms (i.e. liberal bias), human participants exhibited a generally conservative intrinsic bias already with balanced penalties for false alarms and correct rejects. Selectively penalizing false alarms would have aggravated this conservative tendency in humans, hence undermining the cross-species comparison of behavior.

The signal loudness was varied from trial to trial (−30-to-0 dB with respect to RMS noise in mice; (−40-to-5 dB with respect to RMS noise in humans), while the 1 second-mean RMS loudness of the noise was held constant. For the trial containing a signal tone within each mini-block, signal loudness was selected randomly under the constraint that each of six (mice) or five (humans) levels would occur equally often within each session (mice) or block of 60 mini-blocks (humans). The corresponding signal loudness exhibited a robust effect on mean accuracy, with highest accuracy for the loudest signal level: F(5,20) = 23.95, p < 0.001) and F(4,76) = 340.9, p < 0.001), for mouse and human subjects respectively. Human hit rates were almost at ceiling level for the loudest signal (94.7%, ±0.69% s.e.m.), and close to ceiling for the second loudest signal (92.8%, ±0.35% s.e.m.). Because so few errors are not enough to sufficiently constrain the drift diffusion model, we merged the two conditions with the loudest signals.

#### Perceptual (auditory tone-in-noise) yes/no (forced choice) detection task

Each trial consisted of two consecutive intervals (Fig. 3A): (i) the baseline interval (3-4 s uniformly distributed); (ii) the decision interval, the start of which was signaled by the onset of the auditory stimulus and which was terminated by the subject’s response (or after a maximum duration of 2.5 s). The decision interval consisted of only an auditory noise stimulus (McGinley, David, et al., 2015), or a pure sine wave (2 KHz) superimposed onto the noise. In the first experiment, the signal was presented on 50% of trials. Auditory stimuli were presented at the same intensity of 65dB using the same over-ear headphone as in the go/no-go task. In the second experiment, in order to experimentally manipulate perceptual choice bias, the signal was presented on either 30% of trials (“rare” blocks) or 70% of trials (“frequent” blocks) (Fig. 3B). Auditory stimuli were presented at approximately the same signal loudness (65dB) using a Sennheiser HD 660 S over-ear headphone, suppressing ambient noise.

Participants were instructed to report the presence or absence of the signal by pressing one of two response buttons with their left or right index finger, once they felt sufficiently certain (free response paradigm). The mapping between perceptual choice and button press (e.g., “yes” –> right key; “no” –> left key) was counterbalanced across participants. After every 40 trials subjects were informed about their performance. In the second experiment, subjects were explicitly informed about signal probability. The order of signal probability (e.g., first 480 trials –> 30%; last 480 trials –> 70%) was counterbalanced across subjects.

Throughout the experiment, the target signal loudness was fixed at a level that yielded about 75% correct choices in the 50% signal probability condition. Each participant’s individual signal loudness was determined before the main experiment using an adaptive staircase procedure (Quest). For this, we used a two-interval forced choice variant of the tone-in-noise detection yes/no task (one interval, signal-plus-noise; the other, noise), in order to minimize contamination of the staircase by individual bias (generally smaller in two-interval forced choice than yes/no tasks). In the first experiment, the resulting threshold signal loudness produced a mean accuracy of 74.14% correct (±0.75% s.e.m.). In the second experiment, the resulting threshold signal loudness produced a mean accuracy of 84.40% correct (±1.75% s.e.m.) and 83.37% correct (±1.36% s.e.m.) in the rare and frequent conditions, respectively. This increased accuracy was expected given the subjects’ ability to incorporate prior knowledge about signal probability into their decision-making.

#### Memory-based (visual recognition) yes/no (forced choice) decision task

The full experiment consisted of a picture and word encoding task, and a 24 hours-delayed free recall and recognition tests (Fig. 4A) previously described in (Bergt et al., 2018). Here we did not analyze data from the word recognition task because of a modality mismatch: auditory during encoding, visual during recognition. During encoding, 75 neutral and 75 negative greyscale pictures (modified to have the same average luminance) were randomly chosen from the picture pool (Bergt et al., 2018) and presented in randomized order for 3 seconds at the center of the screen, against a grey background that was equiluminant to the pictures. Subjects were instructed to memorize the pictures (intentional encoding) and to evaluate how emotional each picture was on a 4-point scale from 0 (“neutral”) to 3 (“very negative”). During recognition, 24-hours post encoding, subjects saw all pictures that were presented on the first day and an equal number of novel neutral and negative items in randomized order. Subjects were instructed to indicate for each item whether it had been presented the previous day (“yes – old”) or not (“no – new”). For items that were identified as “old”, participants were further asked to rate on a scale from 1 (“not certain”) to 4 (“very certain”) how confident they were that the item was indeed “old”.

### Data acquisition

The mouse pupil data acquisition is described elsewhere (McGinley, David, et al., 2015). The human experiments were conducted in a psychophysics laboratory (go/no-go and yes/no tasks). The left eye’s pupil was tracked at 1000 Hz with an average spatial resolution of 15 to 30 min arc, using an EyeLink 1000 Long Range Mount (SR Research, Osgoode, Ontario, Canada), and it was calibrated once at the start of each block.

### Analysis of task-evoked pupil responses

#### Preprocessing

Periods of blinks and saccades were detected using the manufacturer’s standard algorithms with default settings. The remaining data analyses were performed using custom-made Python scripts. We applied to each pupil timeseries (i) linear interpolation of missing data due to blinks or other reasons (interpolation time window, from 150 ms before until 150 ms after missing data), (ii) low-pass filtering (third-order Butterworth, cut-off: 6 Hz), (iii) for human pupil data, removal of pupil responses to blinks and to saccades, by first estimating these responses by means of deconvolution and then removing them from the pupil time series by means of multiple linear regression (Knapen et al., 2016), and (iv) conversion to units of modulation (percent signal change) around the mean of the pupil time series from each measurement session. We computed the first time derivative of the pupil size, by subtracting the size from adjacent frames, and smoothened the resulting time series with a sliding boxcar window (width, 50 ms).

#### Quantification of task-evoked pupil responses

The auditory yes/no tasks and the yes/no recognition task were analogous in structure to the tasks from our previous pupillometry and decision-making studies (de Gee et al., 2017, 2014). We here computed task-evoked pupil responses time-locked to the behavioral report (button press). We used motor response-locking because motor responses, which occurred in all trials, elicit a transient pupil dilation response (de Gee et al., 2014; Hupé et al., 2009). Thus, locking pupil responses to the motor response balanced those motor components in the pupil responses across trials, eliminating them as a confounding factor for estimates of phasic arousal amplitudes. Specifically, we computed pupil responses as the maximum of the pupil derivative time series (Reimer et al., 2016) in the 500 ms before button press (grey windows in Figs. 3C, S4A, S5A). The resulting pupil bins were associated with different overall pupil response amplitudes across the whole duration of the trial (Figs. S3A, S4B, S5B).

The go/no-go task entailed several deviations from the above task structure that required a different quantification of task-evoked pupil responses. The go/no task had, by design, an imbalance of motor responses between trials ending with different decisions, with no motor response for (implicit) no choices. Thus, no response-locking was possible for no-decisions, leaving stimulus-locking as the only option. In this task, a transient drive of pupil dilation by the motor response (lick or button press) would yield larger pupil responses for go choices (motor movement) than for implicit no-go choices (no motor movement), even without any link between phasic arousal and decision bias. We took two approaches to minimize contamination by this motor imbalance. First, we quantified the single-trial response amplitude as the maximum of the pupil derivative in an early window ranging from the start of the trial-average pupil derivative time course being significantly different from zero up to the first peak (grey windows in Fig. 1D). For the mice, this window ranged from 40–190 after trial onset; for humans, this window ranged from 240–460 ms after trial onset. Second, we excluded decision intervals with a motor response before the end of this window plus a 50 ms buffer (cutoff: 240 ms for mice, 510 ms for humans; Fig. S1D,I). In both species, the resulting pupil derivate defined bins were associated with different overall pupil response amplitudes across the whole duration of the trial (Fig. S1F,K).

For analyses of the go/no-go and yes/no tasks, we used five equally populated bins of task-evoked pupil response amplitudes. We used three bins for the yes/no task with biased environments, because subjects performed substantially fewer trials (see *Subjects*). We used two bins for the recognition task, so that we could perform the individual difference analysis reported in Fig. 4. In the recognition task, we ensured that each pupil bin contained an equal number of neutral and emotional stimuli. In all cases, the results are qualitatively the same when using five equally populated bins of task-evoked pupil response amplitudes.

### Analysis and modeling of choice behavior

In the go/no-go task, the first trial of each mini-block (see *Behavioral tasks*) was excluded from the analyses, because this trial served as a reference and never included the signal (pure sine wave). In the go/no-go and yes/no tasks, reaction time (RT) was defined as the time from stimulus onset until the lick or button press. In the mice go/no-go data set, trials with RTs shorter than 240 ms were excluded from the analyses (see *Quantification of task-evoked pupillary responses* and Fig. S1D); in the human go/no-go data set, trials with RTs shorter than 510 ms were excluded from the analyses (Fig. S1I).

#### Signal-detection theoretic modeling (go/no-go and yes/no tasks)

The signal detection theoretic (SDT) metrics sensitivity (d’) and criterion (c) (Green & Swets, 1966) were computed separately for each of the bins of pupil response size. We estimated d’ as the difference between z-scores of hit rates and false-alarm rates. We estimated criterion by averaging the z-scores of hit rates and false-alarm rates and multiplying the result by -1.

In the go/no-go task, subjects could set only one decision criterion (not to be confused with above-defined c), against which to compare sensory evidence, so as to determine choice. This is because signal loudness was drawn pseudo-randomly on each trial and participants had no way of using separate criteria for different signal strengths. We reconstructed this overall decision criterion (irrespective of signal loudness) and used this as a measure of the overall choice bias, whose dependence on pupil response we then assessed (Fig. 1E). To this end, we used the following approach derived from SDT (Green & Swets, 1966). We computed one false alarm rate (based on the noise trials) and multiple hit rates (one per signal loudness). Based on these we modelled one overall noise distribution (normally distributed with mean=0, sigma=1), and one “composite” signal distribution (Fig. S1C), which was computed as the average across a number of signal distributions separately modelled for each signal loudness (each normally distributed with mean=empirical d’ for that signal loudness, and sigma=1).

We defined the “zero-bias point” *(Z)* as the value for which the noise and composite signal distributions crossed:

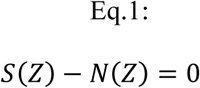

where *S* and *N* are the composite signal and noise distributions, respectively.

The subject’s empirical “choice point” *(C)* was computed as:

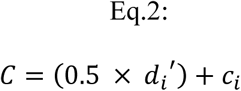

where *di’* and *ci* were a subject’s SDT-sensitivity and SDT-criterion for a given signal loudness, ‘i’. Note that C is the same constant when d’ and criterion are computed for each signal loudness based on the same false alarm rate. Therefore, it does not matter which signal loudness is used to compute the empirical choice point.

Finally, the overall bias measure was then taken as the distance between the subject’s choice point and the zero-bias point:

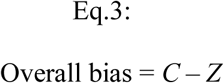

#### Drift diffusion modeling

Data from all tasks were fit with the drift diffusion model, which well captured all features of behavior we assessed. We used the HDDM 0.6.1 package (Wiecki, Sofer, & Frank, 2013) to fit behavioral data from the yes/no and go/no-go tasks. In all datasets, we allowed the following parameters to vary with pupil response-bins: (i) the separation between both bounds (i.e. response caution); (ii) the mean drift rate across trials; (iii) drift bias (an evidence independent constant added to the drift); (iv) the non-decision time (sum of the latencies for sensory encoding and motor execution of the choice). In the datasets using yes/no protocols, we additionally allowed starting point to vary with pupil response bin. In the go/no-go datasets, we allowed non-decision time, drift rate, and drift bias to vary with signal strength (i.e., signal loudness). The specifics of the fitting procedures for the yes/no and go/no-go protocols are described below.

To verify that best-fitting models indeed accounted for the pupil response-dependent changes in behavior, we generated a simulated data set using the fitted drift diffusion model parameters. Separately per subject, we simulated 100000 trials for each pupil bin (and, for the go/no-go data, for each signal loudness), while ensuring that the fraction of signal+noise vs. noise trials matched that of the empirical data; we then computed RT, and signal detection d’ and overall bias (for the go/no-go data sets) or criterion (for the rest) for every bin (as described above).

We used a similar approach to test if, without monitoring task-evoked pupil responses, systematic variations in accumulation bias (drift bias) would appear as random trial-to-trial variability in the accumulation process (drift rate variability) (Fig. 2E). For simplicity, we then pooled across signal loudness and simulated 100000 trials from two conditions that differed according to the fitted drift bias (accumulation bias) estimates in the lowest and highest pupil-defined bin of each individual; drift rate, boundary separation and non-decision time were fixed to the mean across pupil bins of each individual; drift rate variability was fixed to 0.5. We then fitted the drift bias model as described above to the simulated data, and another version of the model in which we fixed drift bias across the two conditions.

#### Yes-no task

We fitted all yes/no datasets using Markov-chain Monte Carlo sampling as implemented in the HDDM toolbox (Wiecki et al., 2013). Fitting the model to RT distributions for the separate responses (termed “stimulus coding” in (Wiecki et al., 2013)) enabled estimating parameters that could have induced biases towards specific choices. Bayesian MCMC generates full posterior distributions over parameter estimates, quantifying not only the most likely parameter value but also the uncertainty associated with that estimate. The hierarchical nature of the model assumes that all observers in a dataset are drawn from a group, with specific group-level prior distributions that are informed by the literature. In practice, this results in more stable parameter estimates for individual subjects, who are constrained by the group-level inference. The hierarchical nature of the model also minimizes risks to overfit the data (Katahira, 2016; Vandekerckhove, Tuerlinckx, & Lee, 2011; Wiecki et al., 2013). Together, this allowed us to simultaneously vary all main parameters with pupil bin: starting point, boundary separation, drift rate, drift bias and non-decision time. We fixed drift rate variability across the pupil-defined bins. We ran 3 separate Markov chains with 12500 samples each. Of those, 2500 were discarded as burn-in. Individual parameter estimates were then estimated from the posterior distributions across the resulting 10000 samples. All group-level chains were visually inspected to ensure convergence. Additionally, we computed the Gelman-Rubin 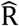 statistic (which compares within-chain and between-chain variance) and checked that all group-level parameters had an 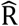 between 0.99-1.01.

#### Go/no-go task

The above described hierarchical Bayesian fitting procedure was not used for the go/no-go tasks because a modified likelihood function was not yet successfully implemented in HDDM. Instead, we fitted the go/no-go data based on RT quantiles, using the so-called G square method (code contributed to the master HDDM repository on Github; https://github.com/hddm-devs/hddm/blob/master/hddm/examples/gonogo_demo.ipynb). The RT distributions for yes choices were represented by the 0.1, 0.3, 0.5, 0.7 and 0.9 quantiles, and, along with the associated response proportions, contributed to G square; a single bin containing the number of no-go choices contributed to G square (Ratcliff et al., 2016). Starting point and drift rate variability were fitted but fixed across the pupil-defined bins. Additionally, drift rate, drift bias and non-decision time varied with signal loudness. The same noise only trials were re-used when fitting the model to each signal loudness.

The absence of no-responses in the go/no-go protocol required fixing one of the two bias parameters (starting point or drift bias) as function of pupil response; leaving both parameters free to vary lead to poor parameter recovery. We fixed starting point based on formal model comparison between a model with pupil-dependent variation of drift bias and starting point: BIC differences ranged from -279.5 to - 137.9 (mean, -235.3: median, -246.6), and from -197.5 to -146.0 (mean, -164.0; median, -162.0) in favor of the model with fixed starting point, for mice and humans respectively. The same was true when ignoring signal loudness: delta BICs ranged from -38.5 to -25.9 (mean, -30.9; median, -29.7), and from -39.8 to -26.7 (mean, -30.9; median, -30.7), for mice and humans respectively.

### Statistical comparisons

We used a mixed linear modeling approach implemented in the *R*-package *lme4* (Bates, Mächler, Bolker, & Walker, 2015) to quantify the dependence of several metrics of overt behavior, or of estimated model parameters (see above), on pupil response. For the go/no-go task, we simultaneously quantified the dependence on signal loudness. Our approach was analogous to sequential polynomial regression analysis (Draper & Smith, 1998), but now performed within a mixed linear modeling framework. In the first step, we fitted three mixed models to test whether pupil responses predominantly exhibited no effect (zero-order polynomial), a monotonic effect (first-order), or a non-monotonic effect (second-order) on the behavioral metric of interest (*y*). The fixed effects were specified as:

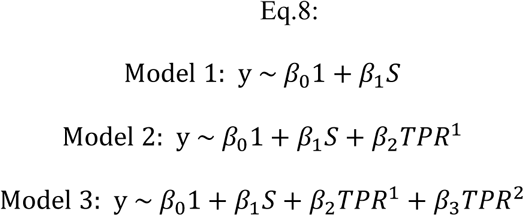

with *β* as regression coefficients, *S* as the signal loudness (for go/no-go task), and TPR as the bin-wise task-evoked pupil response amplitudes. We included the maximal random effects structure justified by the design (Barr, Levy, Scheepers, & Tily, 2013). For data from the go/no-go task, the random effects were specified to accommodate signal loudness coefficient to vary with participant, and the intercept and pupil response coefficients to vary with signal loudness and participant. For data from the yes/no tasks, the random effects were specified to accommodate the intercept and pupil response coefficients to vary with participant. The mixed models were fitted through maximum likelihood estimation. Each model was then sequentially tested in a serial hierarchical analysis, based on chi-squared statistics. This analysis was performed for the complete sample at once, and it tested whether adding the next higher order model yielded a significantly better description of the response than the respective lower order model. We tested models from the zero-order (constant, no effect of pupil response) up to the second-order (quadratic, non-monotonic). In the second step, we refitted the winning model through restricted maximum likelihood estimation, and computed p-values with Satterthwaite’s method implemented in the *R*-package *lmerTest* (Kuznetsova, Brockhoff, & Christensen, 2017).

We used paired-sample t-tests to test for significant differences between the pupil derivative time course and 0, and between pupil response amplitudes for yes versus no choices.

### Data and code sharing

The data are publicly available on [to be filled in upon publication]. Analysis scripts are publicly available on [to be filled in upon publication].

## ACKNOWLEDGMENTS

We thank Daniëlle Rijkmans, Guusje Boomgaard and Christopher David Riddell for help with the data collection for the human auditory detection tasks, Anne Bergt for help with the data collection for the human memory recognition task, and all members of the Donner lab for discussion. This research was supported by the German Research Foundation (DFG, grant numbers: DO 1240/3–1 and SFB 936A7 to THD), European Commission CH2020 7^th^ Framework Programme (Marie Sklodowska-Curie Individual Fellowship: 658581-CODIR, to KT and THD), and the National Institutes of Health (R03DC015618, to MJM).

## AUTHOR CONTRIBUTIONS

JWdG, Conceptualization, Investigation, Formal analysis, Writing—original draft, Writing—review and editing; KT, Conceptualization, Investigation, Formal analysis, Writing—original draft, Writing— review and editing; LS, Conceptualization, Writing—review and editing. AEU, Formal analysis, Writing—review and editing; DAM, Conceptualization, Writing—review and editing; MJM, Conceptualization, Formal analysis, Investigation, Writing—original draft, Writing—review and editing; THD, Conceptualization, Writing—original draft, Writing—review and editing.

## SUPPLEMENTARY FIGURES

**Figure S1.**
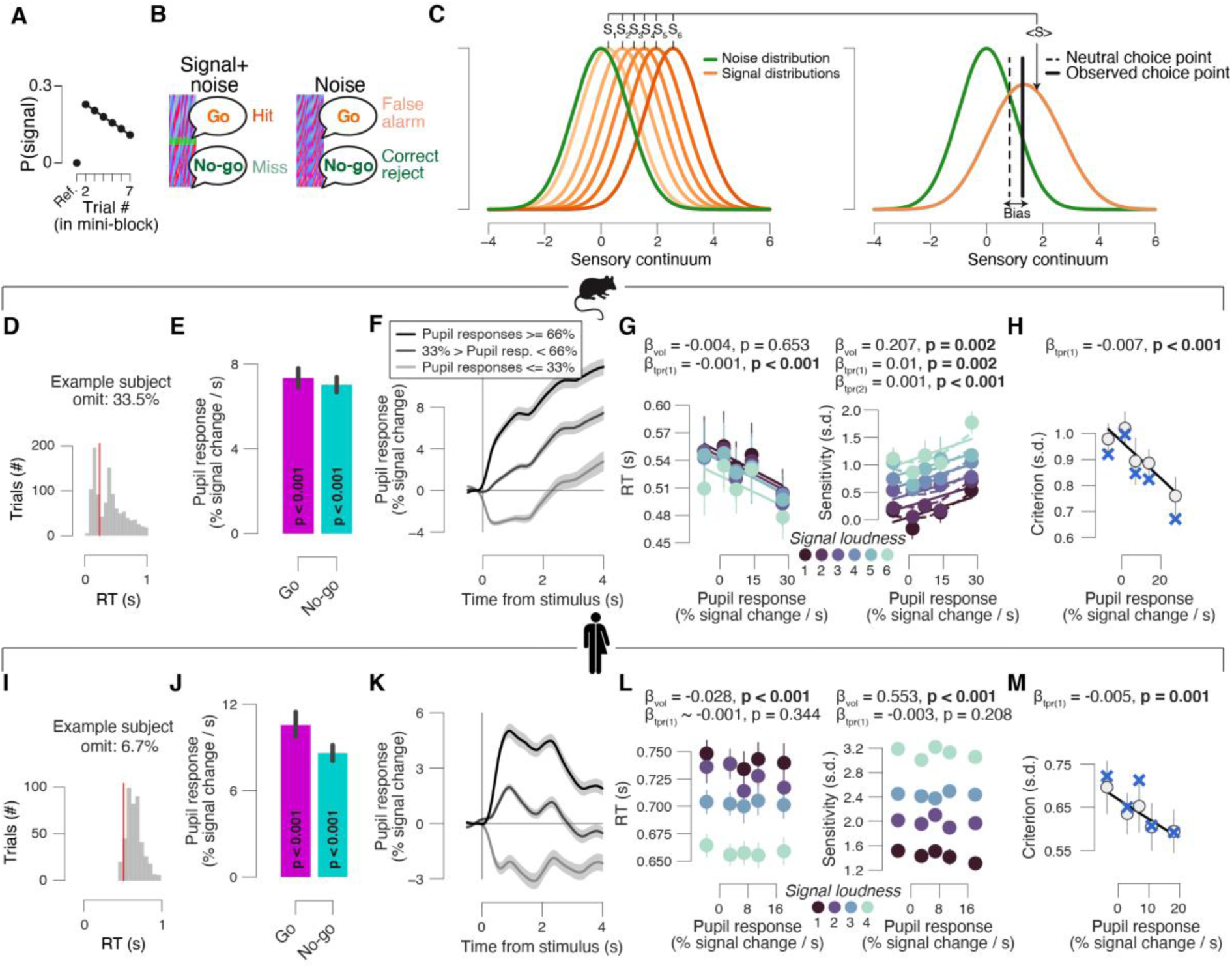
Quantifying pupil responses and behavior in mice and humans. **(A)** Probability of target signal across the sequence of 1–7 trials. Hazard rate of signal occurrence was kept approximately flat. **(B)** The four combinations of stimulus category (signal+noise vs. noise) and behavioral choice (go vs. no-go) yielded the four standard signal detection theory categories. **(C)** Schematic of overall perceptual choice bias measure (Materials and Methods). Per pupil bin we modelled one overall noise distribution (green; normally distributed with mean=0, sigma=1), and one “composite” signal distribution. This composite signal distribution was computed as the average across a number of signal distributions separately modelled for each signal loudness (orange; each normally distributed with mean=empirical d’ for that signal loudness, sigma=1). We defined the “zero-bias point” *(Z)* as the value for which the noise and composite signal distributions cross. The subject’s empirical “choice point” was computed based on the empirical d’ and criterion for any difficulty level (Materials and Methods). The overall bias measure was then taken as the distance between the subject’s choice point and the zero-bias point. **(D)** RT distribution of example subject. Red line, group average latency of the first peak in pupil slope timeseries plus a 50 ms buffer, which was used as a cut-off for excluding trials in order to control for a potential motor confound in our task-evoked pupil response measures (Materials and Methods). Range of omitted trials across all subjects: 29.9%–45.4% (mean, 35.3%; median, 33.5%). **(E)** Task-evoked pupil responses in mice sorted into go and no-go choices (pooled across signal loudness). Stats, paired-samples t-test. **(F)** Overall pupil response time courses in mice for three pupil derivative defined bins (pooled across signal loudness). **(G)** Relationship between median RT (left), perceptual sensitivity (right; quantified by signal detection d’) and pupil response in mice, separately for each signal loudness. Linear fits are plotted wherever the first-order fit was superior to the constant fit (Materials and Methods). Quadratic fits were plotted (dashed lines) wherever the second-order fit was superior to first-order fit. Stats, mixed linear modeling. **(H)** As I, but after computing criterion values separately for all signal strengths and then averaging the resulting criterion values across signal strengths. ‘X’ markers are predictions from best fitting variant of drift diffusion model (Materials and Methods). **(I-M)**, as D-H, but for humans. Range of omitted trials in panel I: 0%–30.7% (mean, 7.0%; median, 4.4%). All panels: group average (N = 5; N = 20); error bars or shading, s.e.m.

**Figure S2.**
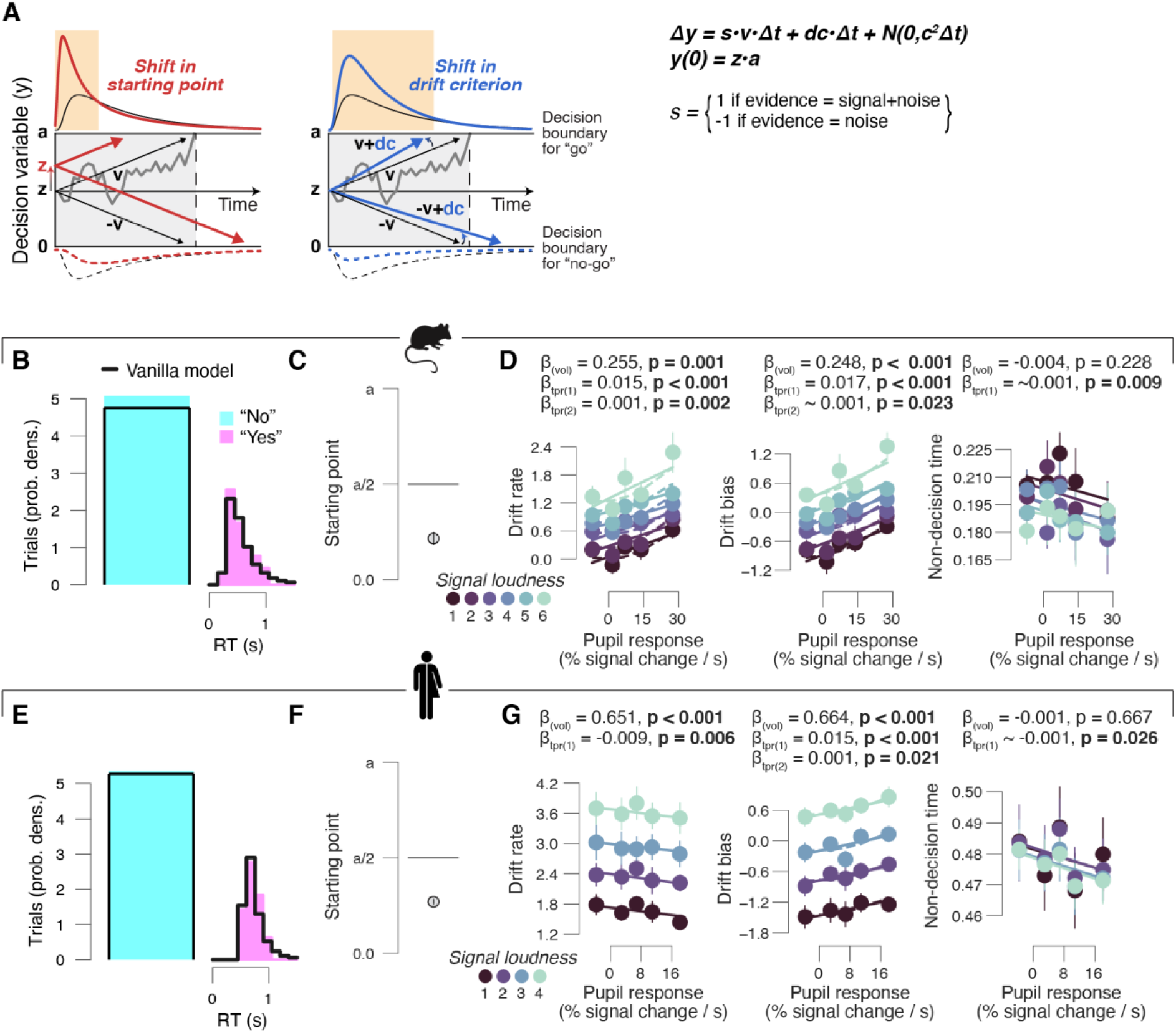
Pupil-dependent changes in computational model parameters during go/no-go task. **(A)** Schematic of drift diffusion model accounting for choices, and their associated RTs (for go trials). Orange windows, RTs for which biased choices are expected under shifts in either “starting point” (*z*; left) or “drift bias” (*dc*; right). Solid (dashed) lines, (implicit) RT distributions. In the equation, *v* is the drift rate (estimated separately for each signal loudness). **(B)** Group average RT distributions, separately for yes and no choices. There were no RTs associated with no choices (no-go); hence, a single bin containing the number of no choices contributed to the model fit (Materials and Methods). Black lines, “vanilla model” fit (parameters boundary separation, drift rate, non-decision time, starting point and drift bias were fixed across signal loudness and pupil bins). **(C)** Starting point estimates of best fitting model (Materials and Methods) in mice expressed as a fraction of the boundary separation (*a*). **(D)** Relationship between drift rate estimates (left), drift bias estimates (right) of best fitting model (Materials and Methods) and pupil responses in mice, separately for each signal loudness. Linear fits are plotted wherever the first-order fit was superior to the constant fit. Quadratic fits were plotted (dashed lines) wherever the second-order fit was superior to first-order fit. Stats, mixed linear modeling. **(E-G)** As B-D, but for humans. All panels: group average (N = 5; N = 20); error bars, s.e.m.

**Figure S3.**
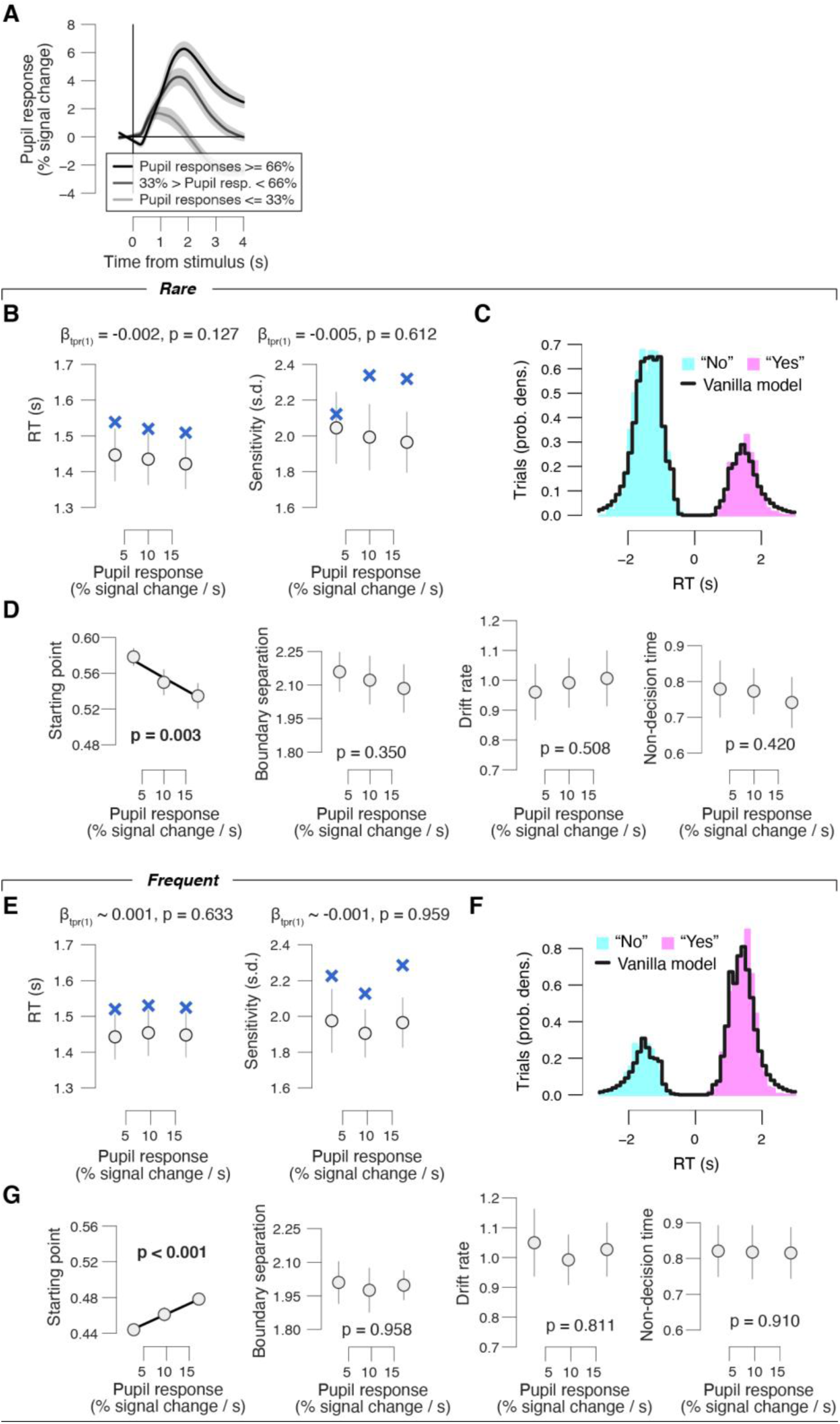
**(A)** Overall pupil response time course for three pupil derivative defined bins. **(B)** Relationship between RT (left), perceptual sensitivity (right) and pupil response in the rare condition. Linear fits are plotted wherever the first-order fit was superior to the constant fit. Quadratic fits were not superior to first-order fits. Stats, mixed-linear modeling. **(C)** Group average RT distributions in the rare condition, separately for yes and no choices. Black lines, “vanilla model” fit (parameters boundary separation, drift rate, non-decision time, starting point and drift bias were fixed across pupil bins). **(D)** As B, but for relationship between drift diffusion model parameters and task-evoked pupil response. **(E-G)** As B-D, for the frequent condition. All panels: group average (N = 14); error bars or shading, s.e.m.

**Figure S4.**
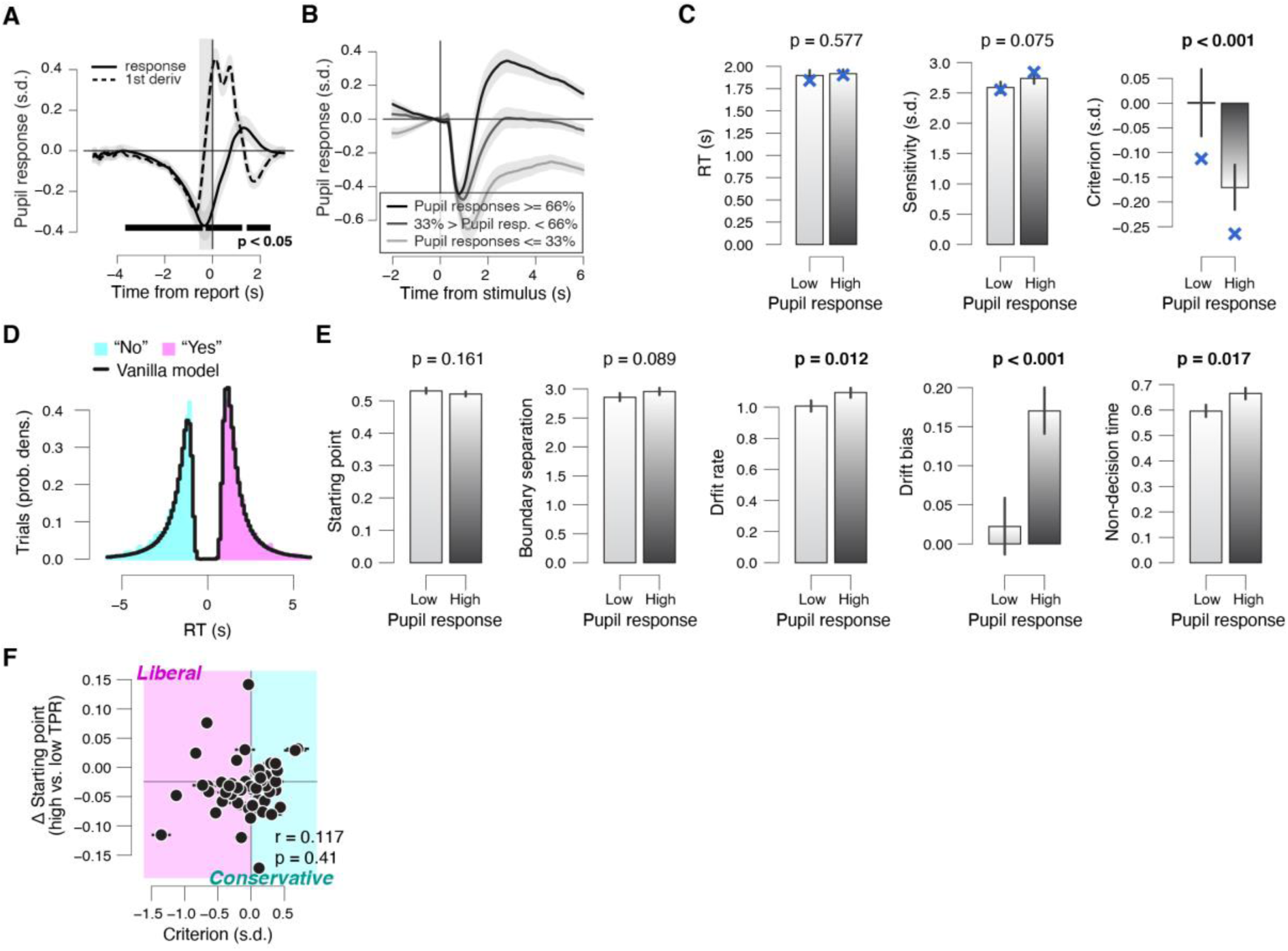
**(A)** Task-evoked pupil response (solid line) and response derivative (dashed line). Grey, interval for task-evoked pupil response measures (Materials and Methods); black bar, significant pupil derivative. Stats, paired-samples t-test. **(B)** Overall pupil response time course for three pupil derivative defined bins. Stats, paired-samples t-test. **(C)** RT (left), sensitivity (right), choice bias, (right) for low and high pupil response bins. **(D)** Group average RT distributions in the conservative condition, separately for yes and no choices. Black lines, “vanilla model” fit (parameters boundary separation, drift rate, non-decision time, starting point and drift bias were fixed across pupil bins). **(E)** As B, but for drift diffusion model parameters. **(F)** Individual pupil predicted shift in starting point, plotted against individual’s overall choice bias. Data points, individual subjects. Stats, Pearson’s correlation. Error bars, 60% confidence intervals (bootstrap). Panels A-E: group average (N = 54); error bars or shading, s.e.m.

**Figure S5.**
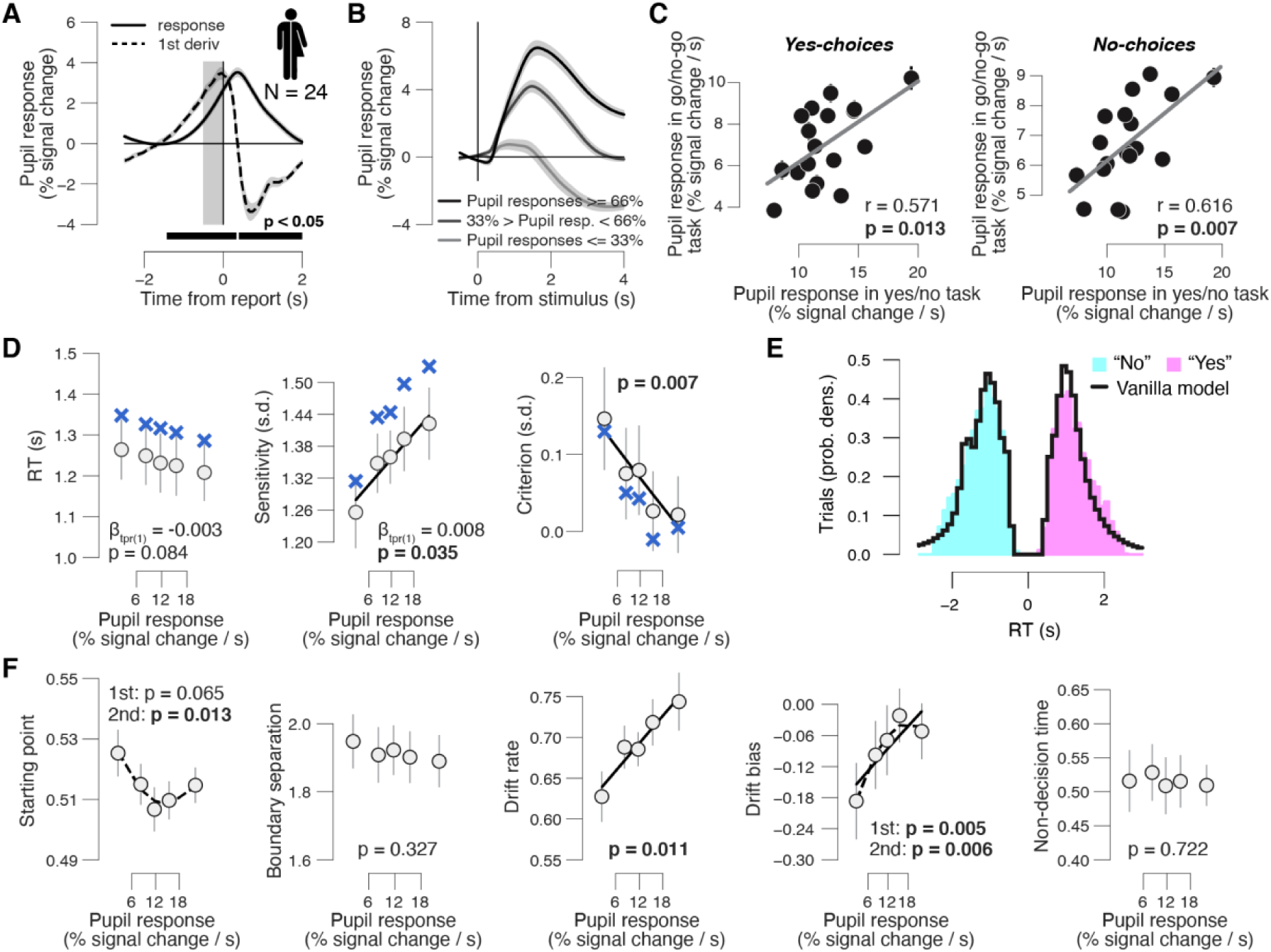
Phasic arousal-related bias suppression does not reflect motor preparation. See Fig. 3A for task-schematic; the results presented here all come from a neutral environment in which the signal occurred on 50% of trials. **(A)** Task-evoked pupil response (solid line) and response derivative (dashed line). Grey, interval for task-evoked pupil response measures (Materials and Methods); black bar, significant pupil derivative. Stats, paired-samples t-test. **(B)** Overall pupil response time course for three pupil derivative defined bins. **(C)** Left: individual task-evoked pupil response amplitude for yes choices in the go/no-go task, plotted against individual pupil response amplitude for yes choices in the yes/no (forced choice) task. Data points, individual subjects. Right: as left, but for no choices. Stats, Pearson’s correlation. A leverage analysis verified that the reported correlations are not driven by outliers. **(D)** Relationship between RT (left), perceptual sensitivity (middle), perceptual choice bias (right) and pupil response. Linear fits were plotted if first-order fit was superior to constant fit; quadratic fits were plotted (dashed lines) wherever the second-order fit was superior to first-order fit. ‘X’ symbols are predictions from the drift diffusion model; stats, mixed-linear modeling. **(E)** Group average RT distributions, separately for yes and no choices. Black lines, “vanilla model” fit (parameters boundary separation, drift rate, non-decision time, starting point and drift bias were fixed across pupil bins). **(F)** As D, but for relationship between drift diffusion model parameters and task-evoked pupil response. Although there was a non-monotonic effect on starting point (p = 0.013), only the pupil-linked changes in drift bias, but not the changes in starting point, strongly correlated with the individual reductions in decision bias as measured by SDT (from panel D) (squared multiple correlation R2 = 0.952; drift bias: beta = -1.01, p < 0.001; starting point: beta = -0.10, p = 0.219). Thus, only the changes of drift bias explained the performance-optimizing reductions in decision bias. All panels: group average (N = 24); error bars or shading, s.e.m.

## REFERENCES

Amaral, D., & Sinnamon, H. (1977). The locus coeruleus: Neurobiology of a central noradrenergic nucleus. Progress in Neurobiology, 9(3), 147–196. https://doi.org/10.1016/0301-0082(77)90016-8

Aston-Jones, G., & Cohen, J. D. (2005). An integrative theory of locus coeruleus-norepinephrine function: Adaptive gain and optimal performance. 28(1), 403–450.

Badre, D., Frank, M. J., & Moore, C. I. (2015). Interactionist Neuroscience. Neuron, 88(5), 855–860.

Barr, D. J., Levy, R., Scheepers, C., & Tily, H. J. (2013). Random effects structure for confirmatory hypothesis testing: Keep it maximal. Journal of Memory and Language, 68(3), 255–278.

Bates, D., Mächler, M., Bolker, B., & Walker, S. (2015). Fitting Linear Mixed-Effects Models Usinglme4. Journal of 1Statistical Software, 67(1).

Beck, J. M., Ma, W. J., Pitkow, X., Latham, P. E., & Pouget, A. (2012). Not noisy, just wrong: The role of suboptimal inference in behavioral variability. Neuron, 74(1), 30–39.

Bergt, A., Urai, A. E., Donner, T. H., & Schwabe, L. (2018). Reading memory formation from the eyes. European Journal of Neuroscience, 47(12), 1525–1533. https://doi.org/10.1111/ejn.13984

Berridge, C. W., & Waterhouse, B. D. (2003). The locus coeruleus-noradrenergic system: Modulation of behavioral state and state-dependent cognitive processes. Brain Research. Brain Research Reviews, 42(1), 33–84.

Bogacz, R., Brown, E., Moehlis, J., Holmes, P., & Cohen, J. D. (2006). The physics of optimal decision making: A formal analysis of models of performance in two-alternative forced-choice tasks. Psychological Review, 113(4), 700–765.

Bouret, S., & Sara, S. J. (2005). Network reset: A simplified overarching theory of locus coeruleus noradrenaline function. Trends in Neurosciences, 28(11), 574–582.

Bowen, H. J., Spaniol, J., Patel, R., & Voss, A. (2016). A Diffusion Model Analysis of Decision Biases Affecting Delayed Recognition of Emotional Stimuli. PLOS ONE, 11(1), e0146769. https://doi.org/10.1371/journal.pone.0146769

Breton-Provencher, V., & Sur, M. (2019). Active control of arousal by a locus coeruleus GABAergic circuit. Nature Neuroscience, 22(2), 218–228. https://doi.org/10.1038/s41593-018-0305-z

Brody, C. D., & Hanks, T. D. (2016). Neural underpinnings of the evidence accumulator. Current Opinion in Neurobiology, 37, 149–157.

Brunton, B. W., Botvinick, M. M., & Brody, C. D. (2013). Rats and humans can optimally accumulate evidence for decision-making. Science (New York, N.Y.), 340(6128), 95–98.

Carandini, M., & Churchland, A. K. (2013). Probing perceptual decisions in rodents. Nature Neuroscience, 16(7), 824–831.

Colizoli, O., de Gee, J. W., Urai, A. E., & Donner, T. H. (2018). Task-evoked pupil responses reflect internal belief states. Scientific Reports, 8(1), 13702. https://doi.org/10.1038/s41598-018-31985-3

Dayan, P., & Yu, A. J. (2006). Phasic norepinephrine: A neural interrupt signal for unexpected events. Network (Bristol, England), 17(4), 335–350.

de Gee, J. W., Colizoli, O., Kloosterman, N. A., Knapen, T., Nieuwenhuis, S., & Donner, T. H. (2017). Dynamic modulation of decision biases by brainstem arousal systems. ELife, 6, 309.

de Gee, J. W., Knapen, T., & Donner, T. H. (2014). Decision-related pupil dilation reflects upcoming choice and individual bias. Proceedings of the National Academy of Sciences of the United States of America, 111(5), E618–25.

Deco, G., Pérez-Sanagustín, M., de Lafuente, V., & Romo, R. (2007). Perceptual detection as a dynamical bistability phenomenon: A neurocomputational correlate of sensation. Proceedings of the National Academy of Sciences, 104(50), 20073–20077.

Donner, T. H., Siegel, M., Fries, P., & Engel, A. K. (2009). Buildup of choice-predictive activity in human motor cortex during perceptual decision making. Current Biology: CB, 19(18), 1581–1585.

Draper, N. R., & Smith, H. (1998). Applied Regression Analysis. Wiley-Interscience.

Drugowitsch, J., Wyart, V., Devauchelle, A.-D., & Koechlin, E. (2016). Computational Precision of Mental Inference as Critical Source of Human Choice Suboptimality. Neuron, 92(6), 1398–1411. https://doi.org/10.1016/j.neuron.2016.11.005

Friston, K. (2010). The free-energy principle: A unified brain theory? Nature Reviews Neuroscience, 11(2), 127–138.

Froemke, R. C. (2015). Plasticity of Cortical Excitatory-Inhibitory Balance. Annual Review of Neuroscience, 38(1), 195–219. https://doi.org/10.1146/annurev-neuro-071714-034002

Gelbard-Sagiv, H., Magidov, E., Sharon, H., Hendler, T., & Nir, Y. (2018). Noradrenaline Modulates Visual Perception and Late Visually Evoked Activity. Current Biology, 28(14), 2239-2249.e6. https://doi.org/10.1016/j.cub.2018.05.051

Gilzenrat, M. S., Nieuwenhuis, S., Jepma, M., & Cohen, J. D. (2010). Pupil diameter tracks changes in control state predicted by the adaptive gain theory of locus coeruleus function. Cognitive, Affective & Behavioral Neuroscience, 10(2), 252–269.

Gold, J. I., & Shadlen, M. N. (2007). The neural basis of decision making. Annual Review of Neuroscience, 30, 535–574.

Green, D. M., & Swets, J. A. (1966). Signal detection theory and psychophysics. 1966. New York.

Harris, K. D., & Thiele, A. (2011). Cortical state and attention. Nature Reviews Neuroscience, 12(9), 509–523.

Hasselmo, M. E. (2006). The Role of Acetylcholine in Learning and Memory. Current Opinion in Neurobiology, 16(6), 710–715. https://doi.org/10.1016/j.conb.2006.09.002

Hsieh, C. Y., Cruikshank, S. J., & Metherate, R. (2000). Differential modulation of auditory thalamocortical and intracortical synaptic transmission by cholinergic agonist. Brain Research, 880(1–2), 51–64.

Hupé, J.-M., Lamirel, C., & Lorenceau, J. (2009). Pupil dynamics during bistable motion perception. Journal of Vision, 9(7), 10.

Jahn, C. I., Gilardeau, S., Varazzani, C., Blain, B., Sallet, J., Walton, M. E., & Bouret, S. (2018). Dual contributions of noradrenaline to behavioural flexibility and motivation. Psychopharmacology, 235(9), 2687–2702. https://doi.org/10.1007/s00213-018-4963-z

Joshi, S., Li, Y., Kalwani, R. M., & Gold, J. I. (2016). Relationships between Pupil Diameter and Neuronal Activity in the Locus Coeruleus, Colliculi, and Cingulate Cortex. Neuron, 89(1), 221–234.

Kane, G. A., Vazey, E. M., Wilson, R. C., Shenhav, A., Daw, N. D., Aston-Jones, G., & Cohen, J. D. (2017). Increased locus coeruleus tonic activity causes disengagement from a patch-foraging task. Cognitive, Affective, & Behavioral Neuroscience, 17(6), 1073–1083. https://doi.org/10.3758/s13415-017-0531-y

Katahira, K. (2016). How hierarchical models improve point estimates of model parameters at the individual level. Journal of Mathematical Psychology, 73, 37–58. https://doi.org/10.1016/j.jmp.2016.03.007

Kepecs, A., Uchida, N., Zariwala, H. A., & Mainen, Z. F. (2008). Neural correlates, computation and behavioural impact of decision confidence. Nature, 455(7210), 227–231.

Kimura, F., Fukuda, M., & Tsumoto, T. (1999). Acetylcholine suppresses the spread of excitation in the visual cortex revealed by optical recording: Possible differential effect depending on the source of input. The European Journal of Neuroscience, 11(10), 3597–3609.

Knapen, T., de Gee, J. W., Brascamp, J., Nuiten, S., Hoppenbrouwers, S., & Theeuwes, J. (2016). Cognitive and Ocular Factors Jointly Determine Pupil Responses under Equiluminance. PLOS ONE, 11(5), e0155574.

Kobayashi, M., Imamura, K., Sugai, T., Onoda, N., Yamamoto, M., Komai, S., & Watanabe, Y. (2000). Selective suppression of horizontal propagation in rat visual cortex by norepinephrine. The European Journal of Neuroscience, 12(1), 264–272.

Krishnamurthy, K., Nassar, M. R., Sarode, S., & Gold, J. I. (2017). Arousal-related adjustments of perceptual biases optimize perception in dynamic environments. Nature Human Behaviour, 1, 0107.

Kuznetsova, A., Brockhoff, P. B., & Christensen, R. H. B. (2017). lmerTest Package: Tests in Linear Mixed Effects Models. Journal of Statistical Software, 82(13).

Lak, A., Nomoto, K., Keramati, M., Sakagami, M., & Kepecs, A. (2017). Midbrain Dopamine Neurons Signal Belief in Choice Accuracy during a Perceptual Decision. Current Biology: CB, 27(6), 821–832.

Larsen, R. S., & Waters, J. (2018). Neuromodulatory Correlates of Pupil Dilation. Frontiers in Neural Circuits, 12.

Lee, S.-H., & Dan, Y. (2012). Neuromodulation of brain states. Neuron, 76(1), 209–222.

Liu, Y., Rodenkirch, C., Moskowitz, N., Schriver, B., & Wang, Q. (2017). Dynamic Lateralization of Pupil Dilation Evoked by Locus Coeruleus Activation Results from Sympathetic, Not Parasympathetic, Contributions. Cell Reports, 20(13), 3099–3112.

Ma, W. J., & Jazayeri, M. (2014). Neural coding of uncertainty and probability. Annual Review of Neuroscience, 37, 205–220.

McGinley, M. J., David, S. V., & McCormick, D. A. (2015). Cortical Membrane Potential Signature of Optimal States for Sensory Signal Detection. Neuron, 87(1), 179–192.

McGinley, M. J., Vinck, M., Reimer, J., Batista-Brito, R., Zagha, E., Cadwell, C. R., … McCormick, D. A. (2015). Waking State: Rapid Variations Modulate Neural and Behavioral Responses. Neuron, 87(6), 1143–1161.

Moran, R. J., Campo, P., Symmonds, M., Stephan, K. E., Dolan, R. J., & Friston, K. J. (2013). Free energy, precision and learning: The role of cholinergic neuromodulation. The Journal of Neuroscience, 33(19), 8227–8236.

Murphy, P. R., Boonstra, E., & Nieuwenhuis, S. (2016). Global gain modulation generates time-dependent urgency during perceptual choice in humans. Nature Communications, 7, 13526. https://doi.org/10.1038/ncomms13526

Murphy, P. R., O’Connell, R. G., O’Sullivan, M., Robertson, I. H., & Balsters, J. H. (2014). Pupil diameter covaries with BOLD activity in human locus coeruleus. Human Brain Mapping, 35(8), 4140–4154.

Najafi, F., & Churchland, A. K. (2018). Perceptual Decision-Making: A Field in the Midst of a Transformation. Neuron, 100(2), 453–462. https://doi.org/10.1016/j.neuron.2018.10.017

Nassar, M. R., Rumsey, K. M., Wilson, R. C., Parikh, K., Heasly, B., & Gold, J. I. (2012). Rational regulation of learning dynamics by pupil-linked arousal systems. Nature Neuroscience, 15(7), 1040–1046.

Parikh, V., Kozak, R., Martinez, V., & Sarter, M. (2007). Prefrontal acetylcholine release controls cue detection on multiple timescales. Neuron, 56(1), 141–154.

Pfeffer, T., Avramiea, A.-E., Nolte, G., Engel, A. K., Linkenkaer-Hansen, K., & Donner, T. H. (2018). Catecholamines alter the intrinsic variability of cortical population activity and perception. PLOS Biology, 16(2), e2003453. https://doi.org/10.1371/journal.pbio.2003453

Pouget, A., Beck, J. M., Ma, W. J., & Latham, P. E. (2013). Probabilistic brains: Knowns and unknowns. Nature Neuroscience, 16(9), 1170–1178.

Pouget, A., Drugowitsch, J., & Kepecs, A. (2016). Confidence and certainty: Distinct probabilistic quantities for different goals. Nature Neuroscience, 19(3), 366–374. https://doi.org/10.1038/nn.4240

Ratcliff, R. (1978). A theory of memory retrieval. Psychological Review, 85(2), 59–108. https://doi.org/10.1037/0033-295X.85.2.59

Ratcliff, R., Huang-Pollock, C., & McKoon, G. (2016). Modeling Individual Differences in the Go/No-Go Task With a Diffusion Model. Decision.

Ratcliff, R., & McKoon, G. (2008). The diffusion decision model: Theory and data for two-choice decision tasks. Neural Computation, 20(4), 873–922.

Reimer, J., Froudarakis, E., Cadwell, C. R., Yatsenko, D., Denfield, G. H., & Tolias, A. S. (2014). Pupil fluctuations track fast switching of cortical states during quiet wakefulness. Neuron, 84(2), 355–362.

Reimer, J., McGinley, M. J., Liu, Y., Rodenkirch, C., Wang, Q., McCormick, D. A., & Tolias, A. S. (2016). Pupil fluctuations track rapid changes in adrenergic and cholinergic activity in cortex. Nature Communications, 7, 13289.

Sara, S. J. (2009). The locus coeruleus and noradrenergic modulation of cognition. Nature Reviews Neuroscience, 10(3), 211–223.

Shadlen, M. N., & Kiani, R. (2013). Decision making as a window on cognition. Neuron, 80(3), 791–806.

Shadlen, M. N., & Shohamy, D. (2016). Decision Making and Sequential Sampling from Memory. Neuron, 90(5), 927–939. https://doi.org/10.1016/j.neuron.2016.04.036

Siegel, M., Engel, A. K., & Donner, T. H. (2011). Cortical network dynamics of perceptual decision-making in the human brain. Frontiers in Human Neuroscience, 5, 21.

Stitt, I., Zhou, Z. C., Radtke-Schuller, S., & Fröhlich, F. (2018). Arousal dependent modulation of thalamo-cortical functional interaction. Nature Communications, 9(1), 2455. https://doi.org/10.1038/s41467-018-04785-6

Tervo, D. G. R., Proskurin, M., Manakov, M., Kabra, M., Vollmer, A., Branson, K., & Karpova, A. Y. (2014). Behavioral Variability through Stochastic Choice and Its Gating by Anterior Cingulate Cortex. Cell, 159(1), 21–32. https://doi.org/10.1016/j.cell.2014.08.037

Urai, A. E., Braun, A., & Donner, T. H. (2017). Pupil-linked arousal is driven by decision uncertainty and alters serial choice bias. Nature Communications, 8, 14637.

van Kempen, J., Loughnane, G. M., Newman, D. P., Kelly, S. P., Thiele, A., O’Connell, R. G., & Bellgrove, M. A. (2019). Behavioural and neural signatures of perceptual decision-making are modulated by pupil-linked arousal. ELife, 8, e42541. https://doi.org/10.7554/eLife.42541

Vandekerckhove, J., Tuerlinckx, F., & Lee, M. D. (2011). Hierarchical diffusion models for two-choice response times. Psychological Methods, 16(1), 44–62. https://doi.org/10.1037/a0021765

Varazzani, C., San-Galli, A., Gilardeau, S., & Bouret, S. (2015). Noradrenaline and Dopamine Neurons in the Reward/Effort Trade-Off: A Direct Electrophysiological Comparison in Behaving Monkeys. Journal of Neuroscience, 35(20), 7866–7877.

Vinck, M., Batista-Brito, R., Knoblich, U., & Cardin, J. A. (2015). Arousal and locomotion make distinct contributions to cortical activity patterns and visual encoding. Neuron, 86(3), 740–754.

Wang, X.-J. (2008). Decision making in recurrent neuronal circuits. Neuron, 60(2), 215–234.

Wiecki, T. V., Sofer, I., & Frank, M. J. (2013). HDDM: Hierarchical Bayesian estimation of the Drift-Diffusion Model in Python. Frontiers in Neuroinformatics, 7, 14.

Wong, K.-F., & Wang, X.-J. (2006). A recurrent network mechanism of time integration in perceptual decisions. The Journal of Neuroscience, 26(4), 1314–1328.

Yerkes, R. M., & Dodson, J. D. (1908). The relation of strength of stimulus to rapidity of habit-formation. Journal of Comparative Neurology and Psychology, 18(5), 459–482.

Zerbi, V., Floriou-Servou, A., Markicevic, M., Vermeiren, Y., Sturman, O., Privitera, M., … Bohacek, J. (2019). Rapid Reconfiguration of the Functional Connectome after Chemogenetic Locus Coeruleus Activation. Neuron, 0(0). https://doi.org/10.1016/j.neuron.2019.05.034

